# Resource allocation in biochemically structured metabolic networks

**DOI:** 10.1101/2024.03.27.586223

**Authors:** Leon Seeger, Fernanda Pinheiro, Michael Lässig

## Abstract

Microbes tune their metabolism to environmental challenges by changing protein expression levels, metabolite concentrations, and reaction rates simultaneously. Here, we establish an analytical model for microbial resource allocation that integrates enzyme biochemistry and the global architecture of metabolic networks. We describe the production of protein biomass from external nutrients in pathways of Michaelis-Menten enzymes and compute the resource allocation that maximizes growth under constraints of mass conservation and metabolite dilution by cell growth. This model predicts generic patterns of growth-dependent microbial resource allocation to proteome and metabolome. In a nutrient-rich medium, optimal protein expression depends primarily on the biochemistry of individual synthesis steps, while metabolite concentrations and fluxes decrease along successive reactions in a metabolic pathway. Under nutrient limitation, individual protein expression levels change linearly with growth rate, the direction of change depending again on the enzyme’s biochemistry. Metabolite levels and fluxes show a stronger, nonlinear decline with growth rate. We identify a simple, metabolite-based regulatory logic by which cells can be tuned to near-optimal growth. Finally, our model predicts evolutionary stable states of metabolic networks, including local biochemical parameters and the global metabolite mass fraction, in tune with empirical data.

## Introduction

Bacteria exhibit remarkable plasticity when reacting to changes in their environment (1, 2). Understanding and harvesting this plasticity promises advances in medicine, synthetic biology, and biotechnology (3–6). Recent improvements in systems-wide metabolic models progress towards this goal (7–10). However, the complexity of metabolic networks and the vast amount of potential model parameters limit a predictive understanding of microbial physiology on the whole-genome level (11–13).

At the same time, simpler empirical patterns emerge from this complexity. Metabolic flux-balance analysis (14), bacterial growth laws governing proteome resource allocation (15), and enzyme-substrate scaling relationships (16) isolate sets of metabolic variables that can be explained without detailed knowledge of the others. Such emergent principles have been used successfully to model the dose-response to ribosome-targeting antimicrobials in diverse environments (17) and to predict dosage-dependent resistance evolution (18). However, dose-response modeling for target enzymes embedded more deeply in the metabolic network than the ribosome (19, 20) remains difficult in general.

In this paper, we integrate enzyme biochemistry and the topology of metabolic pathways into an analytically solvable model of networks with biochemically heterogeneous constituents. The model predicts resource allocation at a multi-omics level including fluxes, the metabolome, and the proteome, as shown schematically in Fig. 1 for a simple chain topology. We consider multi-component metabolic pathways that produce and consume intermediate metabolites, and we compute the state of the cell generating maximal growth under the constraint of mass conservation in a population of replicating cells. This is a step towards an analytical solution of optimal states in large metabolic networks, including realistic non-linear kinetics, which has so far remained elusive (21, 22). The model predicts broad patterns describing how fluxes, protein expression, and steady-state metabolite concentrations depend on an enzyme’s position in the network and on the nutrient environment.

**Figure 1:**
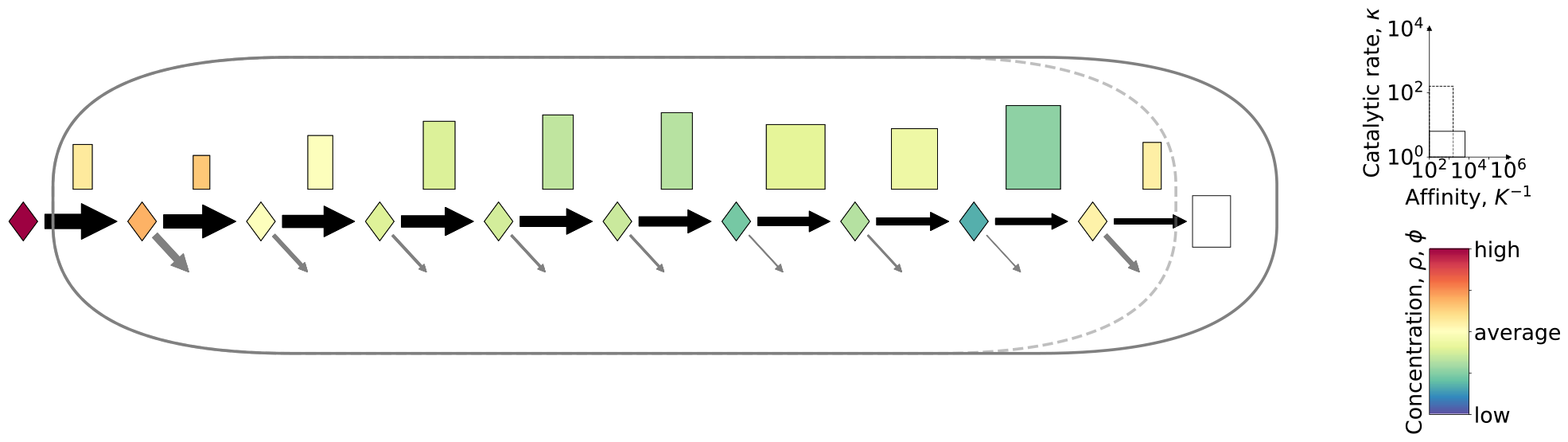
Biochemical model of cell metabolism. A set of reaction pathways processes external nutrients into protein production, leading to cell growth. A simple network is a metabolic chain of 𝓁 enzymes (boxes) and 𝓁 metabolites (diamonds), shown here for 𝓁 = 10. Each reaction step is characterized by Michaelis-Menten kinetics with catalytic capacity 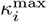 (box height) and substrate binding affinity 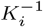 (box width) (*i* = 1, …, 𝓁). The first metabolite is an external nutrient of mass density *ρ*_1_, the output of the network is protein production at rate λ. The model developed in this paper predicts growth-optimal, balanced growth states in nutrient-rich environments and under nutrient limitation. Such states are characterized by steady-state metabolite concentrations, *ρ*_*i*_, and protein expression levels, ϕ_*i*_ (color shading). The network produces a decreasing sequence of mass fluxes, *j*_*i*_ (black arrows), with loss of metabolites by growth (gray arrows).

A key network parameter, called the metabolite cost, summarizes the effects of nutrient level and upstream metabolism on a given metabolite-enzyme pair. This constraint combines with local metabolic reaction kinetics to set the resource allocation to individual metabolites and enzymes. Under nutrient restriction, an enzyme is linearly up- or down-regulated with decreasing growth rate. The direction of change depends on a local biochemical parameter negatively correlated with the enzyme’s target affinity and positively correlated with its catalytic rate. This pattern explains the heterogeneity of single-enzyme growth responses found in proteomics studies of microbial growth (23–25).

An integrative metabolic model of enzymes and metabolites is key to understand how regulation and evolution shape systems metabolism. Here we introduce a simple regulatory scheme that suggests how cells can be tuned to near-optimal growth by using metabolites as transcriptional signals for protein expression. On longer time scales, evolution alters network biochemistry. Specifically, we show that physiological values of enzyme kinetics emerge from an evolutionary equilibrium. This, in turn, sets the global mass ratio of metabolome and proteome in the cell.

The text is organized as follows. We first introduce the model framework and derive local optimality conditions that determine cellular resource allocation to a given metabolite-enzyme pair. We then solve these equations, first in a nutrient-rich environment, and discuss the resulting system-wide patterns of metabolite levels and protein expression. Next, we analyze the metabolic response to nutrient limitation, explore more complex metabolic networks, and compare our findings to published experimental data (16, 24). Finally, we discuss the implications of the model for regulation and evolution of metabolic networks.

## Metabolic model

### Networks of metabolites and enzymes

We consider a metabolic network of 𝓁 enzymes. Each enzyme catalyzes a reaction according to Michaelis-Menten kinetics: in a cell of volume *V*, the enzyme species present at mass *M*_*i*_ acts on a metabolite present at mass concentration *ρ*_*i*_ with catalytic rate 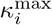 and substrate binding affinity 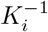, generating a mass flux

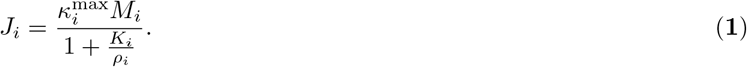

The metabolite mass flux per unit of protein mass, 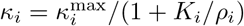 is the product of 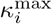 and the saturation level of the enzyme with its substrate. In line with empirical observations (1, 26–28), we assume that the total intracellular protein concentration is constant (15); the cell volume is therefore proportional to the total protein biomass 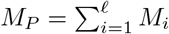. Accordingly, we write concentrations as mass fractions, *ρ*_*i*_ = *R*_*i*_/*M*_*P*_, *ϕ*_*i*_ = *M*_*i*_/*M*_*P*_, and we use mass-normalized fluxes, *j*_*i*_ = *J*_*i*_/*M*_*P*_, and dimensionless affinities, 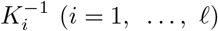. Details are given in Supporting Information (SI). These affinities set the ratio of intracellular metabolite biomass to protein biomass, 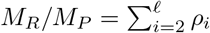 fundamental global observable of cell metabolism. The metabolic state of the system, denoted by the shorthand 𝒮 ≡ (*ρ, ϕ*), changes in response to the environment. The set of kinetic constants, (*κ*^max^,*K*), is fixed for a given organism but changes by evolution on longer time scales.

We start with the simplest case of a metabolic network, which consists of a single linear pathway where the metabolite produced by one enzyme is the substrate of the next (Fig. 1). Overall, the network converts a single environmental nutrient, present at concentration *ρ*_1_, into protein biomass with a rate

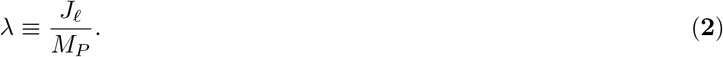

This defines the growth rate of the cell population. In the case of a single pathway, mass conservation on all intracellular metabolites takes the form

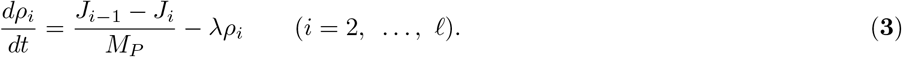

We consider balanced exponential growth states (29) (*ρ*_*i*_ = const. for *i* = 2, …, *𝓁*) in a time-independent environment *ρ*_1_. Importantly, these states maintain a flux mismatch 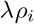 between metabolite in- and outflux at each metabolic step, reflecting the dilution of metabolites by growth (8, 22, 30–33). With the boundary condition *j*_*𝓁*_ = *λ*, we obtain a decreasing cascade of steady-state mass fluxes,

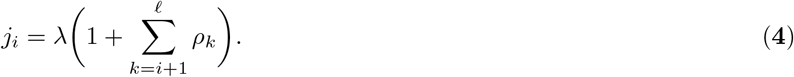

### Optimal balanced growth states

In the following, we consider metabolic states that maximize growth rate under given nutrient conditions, 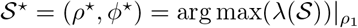. Under laboratory conditions, growth rate is often closely related to fitness (34), and bacteria allocate their resources close to the growth-optimal allocation (35). This motivates our assumptions on stationarity and growth-maximization as a reasonable approximation of more complex growth cycles and evolutionary objectives (5).

Finding optimal balanced growth states analytically is challenging because the kinetic rate law is nonlinear and there is a global feedback loop imposed by growth dilution: the dilution rate in each step is set by growth rate, which itself depends on the output of the pathway. Environmental conditions and cell growth thus impose constraints from opposite ends of the metabolic cascade. Partially numerical solutions (30), specific properties of optimal solutions (8, 22), as well as an analytical solution of a pathway with two enzymes (33) were studied previously. Here, starting from the two-enzyme system, we iteratively construct optimal balanced growth states of longer pathways by adding a substrate-enzyme pair to the front end (SI, Fig. S1).

By equation [**2**], maximal growth is achieved if the protein mass necessary to produce a given protein production flux *J*_*𝓁*_ is minimal (36). We use the Michaelis-Menten relation [**1**] and the continuity equation [**4**] for each reaction to express the enzyme masses *M*_*k*_ in an arbitrary stationary state in terms of the nutrient level *ρ*_1_, the current vector *J* = (*J*_1_, …, *J*_*𝓁*_), and the set of kinetic constants (*κ*^max^,*K*). Thus, we can formulate the maximum-growth condition in differential form,

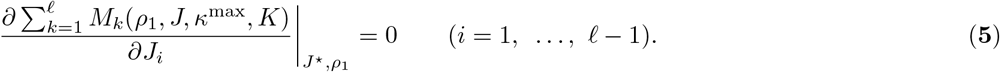

When a single flux *J*_*i*_ is varied, only the metabolite concentrations *ρ*_*i*_, *ρ*_*i*+1_ and the enzyme masses *M*_*i*_, *M*_*i*+1_ change (Fig. S1C). Inserting the Michaelis-Menten relationship [**1**] for *M*_*i*_ and *M*_*i*+1_, as well as equation [**4**] for *ρ*_*i*_ and *ρ*_*i*+1_, then yields a recursive relation for growth-optimal fluxes,

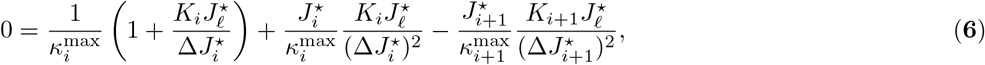

where 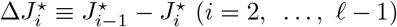. The initial condition 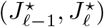 parameterizes the growth rate λ and the (a priori unknown) protein mass *M*_*P*_. Solving this recursion produces the chain of metabolic fluxes, 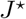 by equations [**1**]–[**4**], we then obtain *M*_*P*_ and the mass-normalized metabolic state *S*^⋆^ = (*ρ* ^⋆^, *ϕ*^⋆^). To study environmental perturbations, we can alternatively express *S*^⋆^ as a function of the nutrient level *ρ*_1_ or the growth rate λ as the independent control parameter. Details of the solution procedure are given in SI.

## Results

### Local optimality and metabolite cost

We now study local properties of optimal balanced growth states in more detail. To this end, we consider a variation around the growth-optimal state that alters a single metabolite concentration *ρ*_*i*_, whereas all other metabolite concentrations and the protein production flux *J*_*𝓁*_ = *λM*_*P*_ remain constant (Fig. S1D). The optimality criterion (**5**) takes the form

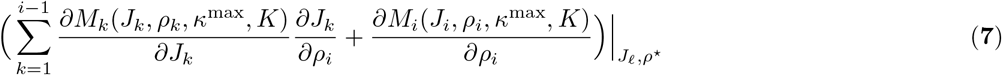

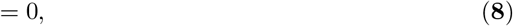

where ∂*M*_*k*_/*∂J*_*k*_ = 1/*κ*_*k*_ by the Michaelis-Menten relation [**1**] and *∂J*_*k*_/*∂ρ*_*i*_ = *λM*_*P*_ by mass conservation [**4**]. Thus, a small increase in concentration of the focal metabolite, *δρ*_*i*_ *>* 0, is generated by an expression decrease of its cognate enzyme, *δM*_*i*_ *<* 0, and an increase of all upstream enzymes, *δM*_*k*_ *>* 0 for *k < i*. All metabolic currents *J*_*i*_, *J*_*i*+1_, …, *J*_*𝓁*_ and all masses *M*_*i*+1_, …, *M*_*𝓁*_ remain constant to first order. We can rewrite this relation for intensive variables,

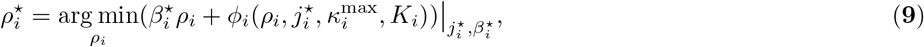

where the *metabolite cost* parameter 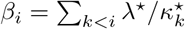 accounts for the upstream proteome resources required to sustain an increased focal metabolite concentration against metabolite dilution in nutrient-rich medium. A more general form of this relation applies when the external nutrient becomes growth-limiting (*ρ*_1_ ≲ *K*_1_) and the nutrient uptake enzyme is constrained 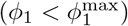. This constraint, the biological rationale of which is discussed below, curbs all downstream metabolite concentrations, *ρ*_2_, …, *ρ*_𝓁_. It can be described by an additional metabolite cost 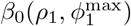 that becomes relevant under nutrient restriction and is negligible in the nutrient-rich regime (SI). Together, we obtain the total metabolite cost

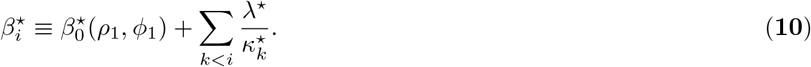

This cost summarizes the influence of the global metabolic network and the nutrient environment on the local balance of substrate and enzyme masses.

### Metabolite-enzyme relations

Combining the optimality condition [**9, 10**] with the kinetic rate law [**1**], we obtain remarkably simple constitutive relations for cellular resource allocation,

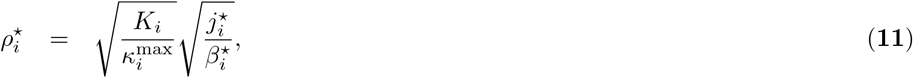

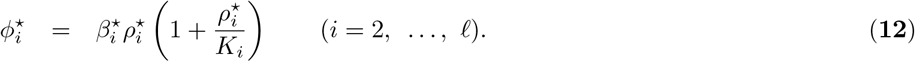

The first relation determines *ρ*_*i*_ as a function of the local kinetic parameters *K*_*i*_, 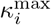 and the network-dependent variables *j*_*i*_, β_*i*_. The second relation links enzyme and metabolite mass fractions. Fig. 2 summarizes the central results of this model; derivations and more detailed analysis will be given in the remainder of this section. In maximum-growth states (*ρ*^⋆^, *ϕ*^⋆^), resource allocation to individual metabolite-enzyme pairs depends on three main factors: network position, nutrient level, and the biochemistry of local metabolic steps (arrows in Fig. 2). This pattern is organized by two key parameters: the metabolite cost β_*i*_ and the growth response parameter 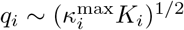, a summary variable of local reaction kinetics (grid lines in Fig. 2). The cost parameter β_*i*_ encodes information on network position and on the environment. Increasing cost generates reduced metabolite levels *ρ*_*i*_ and a weaker, inhomogeneous response of enzymes: high-*q* proteins increase and low-*q* proteins decrease in their expression level *ϕ*_*i*_. Variation of the growth response parameter *q* induces a crossover from a linear enzyme-metabolite relation at low saturation (*ϕ*_*i*_ ∼ *ρ*_*i*_ for *ρ*_*i*_/*K*_*i*_ ≲ 1) to a quadratic relation at high saturation (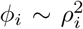 for *ρ*_*i*_/*K*_*i*_ ≳ 1), as given by equation [**12**]. A similar scaling relation has been derived by Dourado et al. (16) from a mass minimization condition for an individual metabolite-enzyme pair. The network relation [**12**] differs in two aspects: the metabolite density is independently set by equation [**11**] and is weighed by the cost parameter *β*_*i*_.

**Figure 2:**
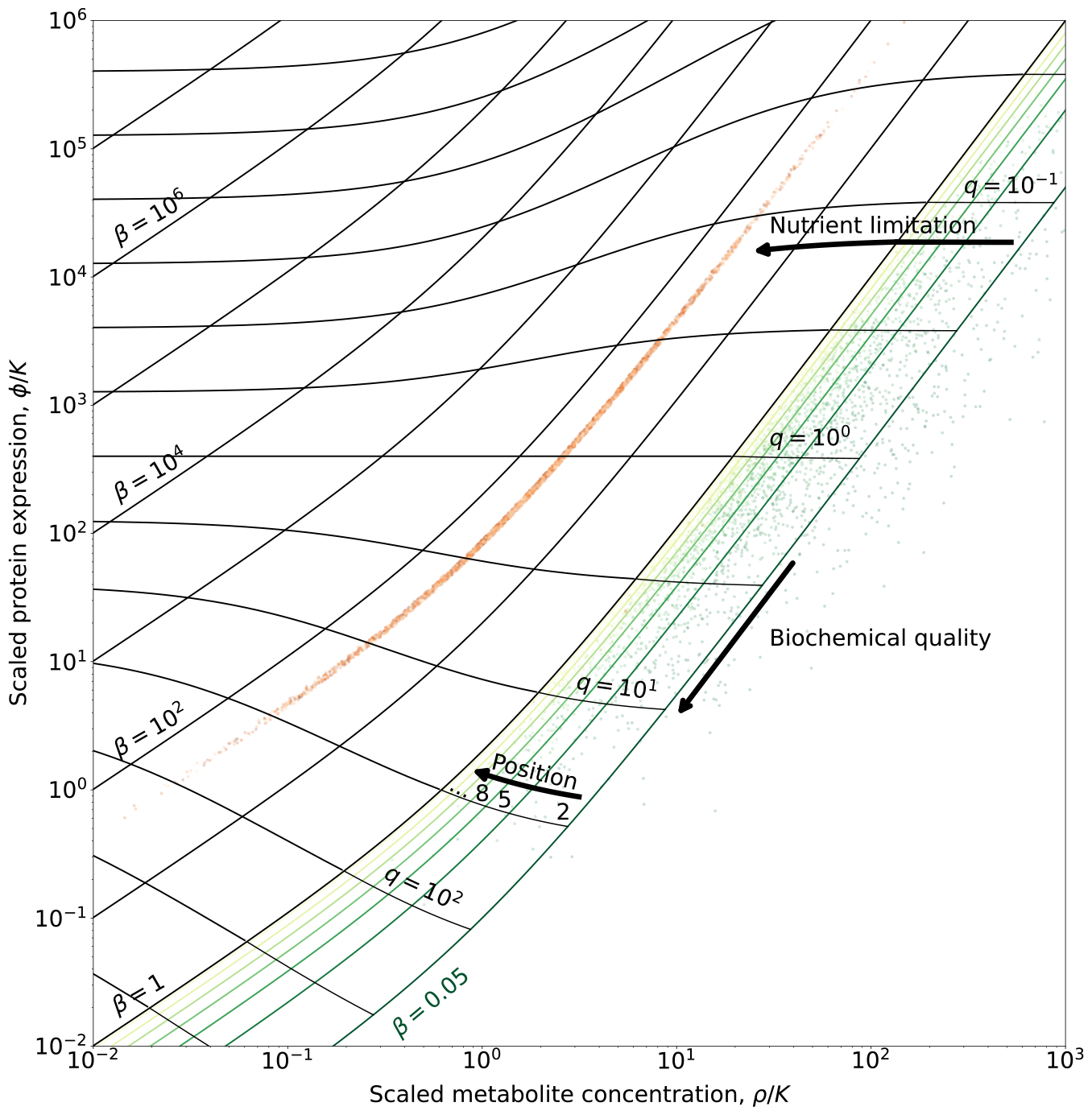
Resource allocation to metabolites and enzymes in metabolic networks. Points show affinity-scaled mass fractions of individual metaboliteenzyme pairs, 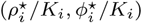, in optimal balanced states of metabolic networks at high nutrient supply (green, 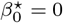) and under nutrient limitation (orange,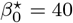). These levels are determined by two key parameters (solid lines): the metabolite cost β_*i*_ and the growth response parameter 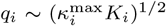, which depends on the kinetic coeffcients of the metabolite-enzyme pair (see text for precise definitions). At high nutrient supply, the cost depends on the network position *i*, causing higher levels of upstream metabolites (darker green shading), while protein expression depends predominantly on local biochemistry; see also Fig. 3A, B. Nutrient limitation induces a global increase of metabolic costs, generating a strong decrease of metabolite levels and a weaker, inhomogeneous response of protein expression: high-*q* proteins are up-regulated, low-q proteins are down-regulated with decreasing nutrient supply; see also Fig. 4. Model parameters: chain length 𝓁 = 20; kinetic parameters are log-normal random variables, 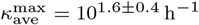, *K*_ave_ = 10^*-*4.2*±*1.1^, resulting in networks with dilution strength ϵ = 0.06±0.04,, as given by equation [**17**]. Grid lines and nutrient limitation (orange points) are shown for specific realizations with ϵ= 0.05 and 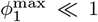. In these networks, 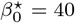 corresponds to a growth reduction by 50 %.

### Resource allocation at high nutrient supply

We now show that the constitutive relations [**12**] and [**11**], together with the continuity relation [**4**] and the cost function [**10**], determine cellular states of maximum growth. We start with cells in a nutrient-rich environment, which grow at a rate close to their physiological bound. In this growth regime, we find a self-consistent solution

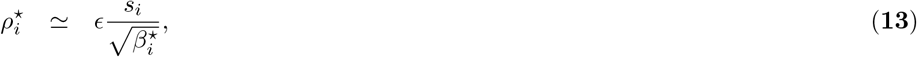

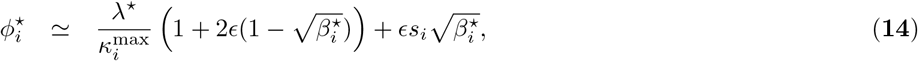

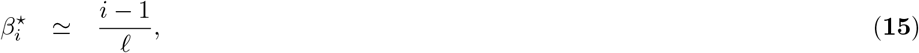

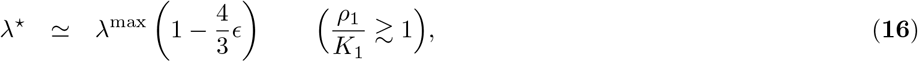

where 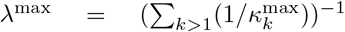 is the maximum growth rate in the absence of metabolite dilution,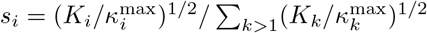 depends on the enzyme specificity 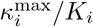, and

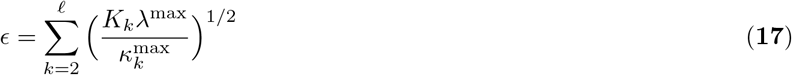

is a dimensionless biochemical summary variable, which we term *dilution strength*. To derive these relations, consider first the case ϵ = 0, which reproduces the result of standard flux balance analysis: fluxes are conserved throughout the network (*j*_*i*_ = *λ*), all enzymes run at full saturation with rates 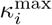 and optimal expression levels proportional to 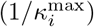, and metabolites are neglected (*ρ*_*i*_ = 0). The effects of metabolite dilution can be included by a power series expansion in the Parameter ϵ. Equations [**13**, **14**] reproduce the constitutive relations [**11, 12**] to first order in *ϵ*. In the following steps, we use a mean-field approximation, replacing the heterogeneous kinetic coefficients 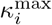, *K*_*i*_ by position-independent values 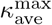, *K*_ave_. To evaluate the metabolite cost, we approximate [**10**] as 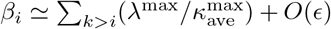, which leads to equation [**15**]. In the same way, we approximate the continuity equation [**4**] to evaluate the effect of flux degradation on protein expression,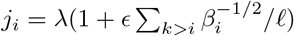, which enters equation [**14**]. Finally, the first-order correction to the growth rate, equation [**16**],is obtained from the normalization condition ∑*kϕk* = 1. This relation predicts the degradation of growth by metabolite dilution in typical metabolic networks.

Physiological values of the parameter *ϵ* can be inferred from empirical data. Metabolic networks have typical catalytic rates *κ*^max^ ≈10^1.6*±*0.4^ h^*-*1^ and Michaelis constants *K ≈*10^*-*4.2*±*1.1^ in dimensionless units of mass fractions (37). Typical pathways of protein biosynthesis from glucose via a representative amino acid have length 𝓁 *≈* 20 (SI). In mean-field approximation, equation [**17**] then yields typical values *ϵ* ≈ 0.05 of the dilution strength, justifying the first-order calculus in equations [**13**] – [**16**]. We can also use these physiological network data to test the accuracy of the mean-field approximation. We construct a set of metabolic chains with kinetic parameters 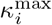, *K*_*i*_ (*i* = 1, …, 20) independently drawn from log-normal distributions centered at the corresponding physiological average values (SI). Consistently, these chains have growth rates of order 1 h^*-*1^, and match other physiological cell properties (Fig. S2). For each chain, we evaluate maximum-growth metabolic state *S*^⋆^ = (*ρ*^⋆^, *ϕ*^⋆^) by numerical solution of equation [**6**] and compare with equations [**13** – **16**]. Remarkably, the analytical form reproduces individual metabolite and enzyme levels of heterogeneous chains, as well as chain-specific growth rates, in good approximation (Fig. 3, Fig. S2E).

**Figure 3:**
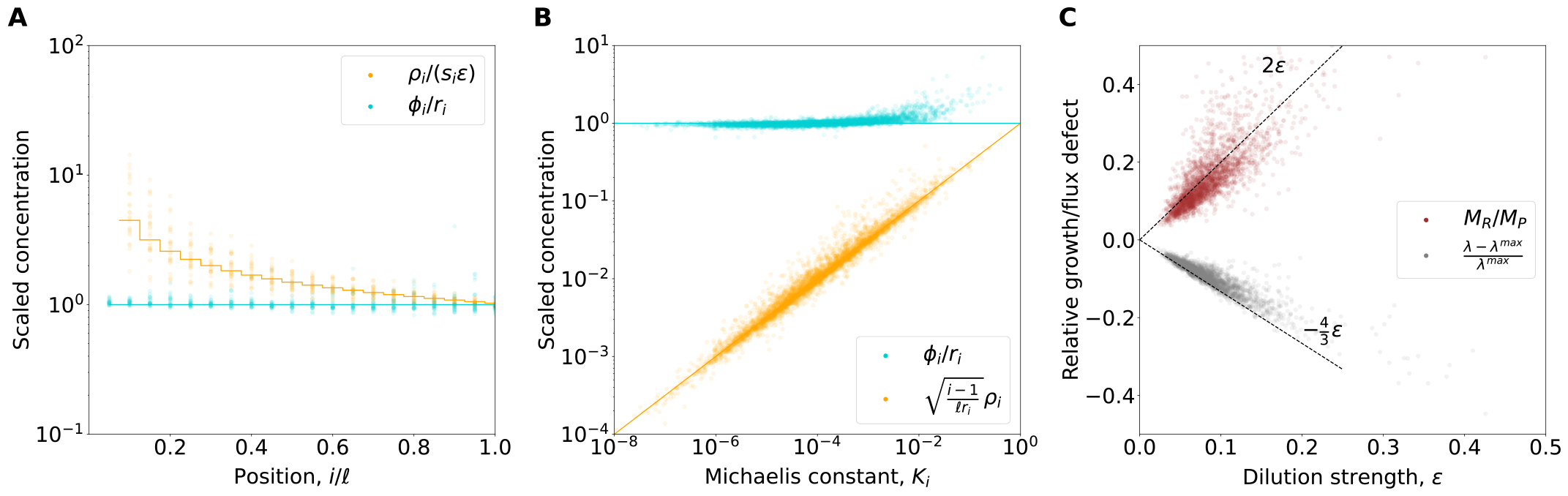
Metabolite effects at high nutrient supply. (*A*) Specificity-scaled metabolite and enzyme catalytic rate-scaled protein levels, *ρ*_*i*_/s_*i*_ and ϕ_*i*_/*r*_*i*_, as functions of relative network position, i/𝓁 (with 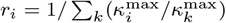). Points are individual enzyme-substrate pairs in physiologically parameterized cell models in a nutrient-rich environment. Lines are analytical predictions of equations [**13** – **17**]. (*B*) Position- and catalytic rate-scaled metabolite and protein levels, *ρ*_*i*_(*β*_*i*_/*r*_*i*_)^1/2^ and ϕ_*i*_/*r*_*i*_, as functions of inverse substrate affinity, *K*_*i*_. (*C*) Growth rate degradation, λ/λ^max^, and metabolome-proteome mass ratio, *M*_*R*_/*M*_*P*_, as functions of the dilution strength ϵ. Points are complete, physiologically parameterized cell models in a nutrient-rich environment. Stochastic parameters as in Fig. 2.

Together, our model predicts a consistent pattern of metabolic resource allocation in nutrient-rich media. Metabolite levels strongly depend on network position and on enzyme specificity, 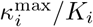 metabolites in upstream or less specific reactions have systematically higher concentrations than those in downstream or more specific reactions (Fig. 3A, B). Enzymes show a weaker dependence on these parameters; their expression levels are predominantly determined by the catalytic rate, 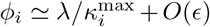. The corresponding affinity-scaled levels are predominantly in the high-saturation regime, *ρ*_*i*_/*K*_*i*_, *ϕ*_*i*_/*K*_*i*_ ≳ 1 (Fig. 2), consistent with empirical data (16, 38).

### Metabolome-proteome mass ratio

Our model predicts the relative abundance of metabolites and proteins in a metabolic network as a function of its reaction kinetics. We sum up the metabolite densities *ρ*_*i*_, using equation [**13**] in mean-field approximation 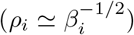, to obtain the metabolome-proteome mass ratio in typical metabolic networks at high nutrient supply,

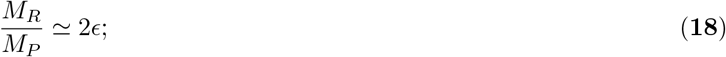

with correction terms of order *O*(*ϵ*^2^, *ϵ*/ *𝓁*) (SI). The resulting numerical value, *M*_*R*_/*M*_*P*_ *∼*10^*-*1^, is in line with empirical data (38, 39), providing a further consistency check for the model. Equations [**16**, **17**] measure the growth cost of metabolites at high nutrient supply; this cost is caused by dilution and increases with the network size *ℓ*. Below, we show that physiological values of *M*_*R*_/*M*_*P*_ are set by the global evolution of network biochemistry.

### Nutrient limitation

Cells respond to a reduced density of external nutrients by up-regulation of the cognate uptake machinery. In our simple metabolic chain, we model this response as a nutrient-dependent expression of the first enzyme, 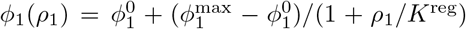, which varies between a basal level 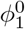 at high nutrient concentration *ρ*_1_ and a maximal level 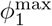 at low concentration (Fig. S3A). The upper bound 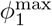 can be generated by different biological mechanisms, including spatial constraints on the cell membrane, diffusive limits on nutrient uptake at low supply (40), and a maximum fold change achievable by regulation. Such limited overexpression of the uptake machinery has been observed for glucose limitation in chemostat experiments (41). In turn, this generates a constrained optimal metabolic state,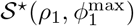. In this state, the growth rate is limited by the uptake step,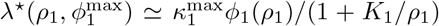. Nutrient limitation significantly affects growth in the regime *ρ*_1_/*K*_1_ ≲ 1, corresponding to 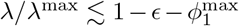 (SI, Fig. S2F). In the following, we use *λ* as an independent control parameter. Under strong nutrient limitation, the constitutive relations [**11, 12**] take the form,

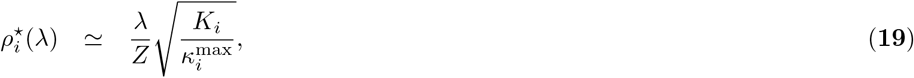

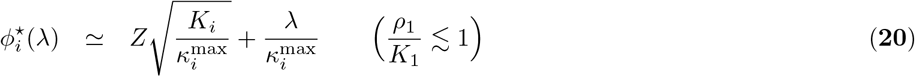

for *i* = 2, …, *𝓁*, where the normalization factor

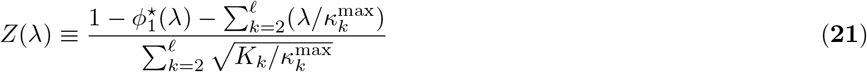

is given by the condition ∑_*k*_ *ϕ*_*k*_ = 1. Here we have omitted the flux degradation by metabolite dilution (*j*_*i*_ *≃ λ*), which becomes small at low growth rates. As shown by comparison of equations [**19**, **20**] and [**10** – **12**], nutrient limitation generates an overall metabolite cost (*β*_0_ = *Z*^2^/*λ*. This cost is numerically larger than the network-dependent terms 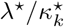, which are therefore neglected (*β*_*i*_ *≃ β*_0_). For simplicity, we also assume that the uptake proteins remain a small fraction of the total proteome 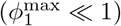, which simplifies equation [**21**] to 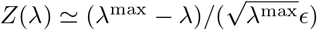.

The solution [**19** – **21**] shows two salient features of metabolic response to nutrient limitation. Given that *Z* is a linear function of λ, metabolite levels increase in a strongly nonlinear way with growth rate,

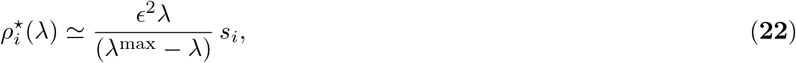

With 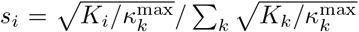. This equation is correct up to position-dependent pre-factors of order 1 relevant at high growth, and differences in *ϕ*_1_ (Fig. 4B). The global metabolome-proteome mass ratio, *M*_*R*_/*M*_*P*_, depends on growth in the same way. In contrast, protein expression levels show a linear response pattern,

**Figure 4:**
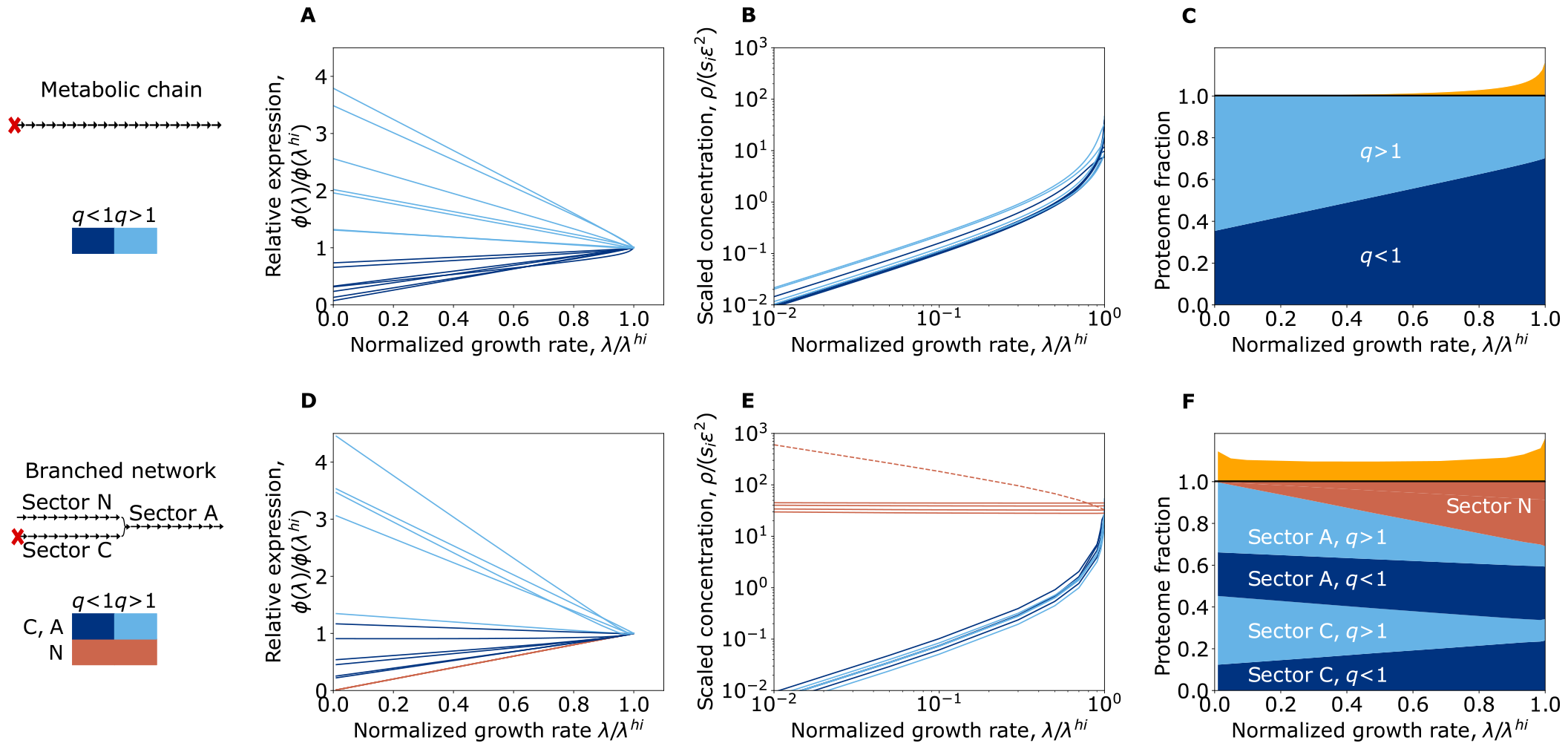
Metabolic response to nutrient limitation. (*A - C*) Metabolic chain. (*A*) Reduction of protein expression levels, 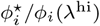, as functions of the growth rate, for two enzyme classes: rate-optimized (*q* > 1) and specificity-optimized (*q* < 1). Here λ^hi^ is the maximum physiological growth rate at high nutrient supply, as given by equation [**16**]. Lines represent randomly drawn enzymes from the ensemble of physiologically parametrized networks (see text; nutrient uptake enzymes were excluded from the analysis). (*B*) Reduction of metabolite levels, 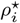, as functions of the growth rate. (*C*) Ensemble averages of global response of proteome and metabolome. The metabolite mass fraction is shown in units of the proteome mass in orange. (*D - F*) Three-sector system. (*D*) Growth-dependent protein levels, 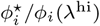, by growth response class and by sector. (*E*) Growth-dependent metabolite levels, 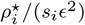, by sector. (*F*) Global response of proteome and metabolome. Chain length: 𝓁 = 20, 𝓁_*A*_ = 𝓁_*C*_ = 𝓁_*N*_ = 10. Model parameters as in Fig. 2 for chains with 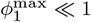.

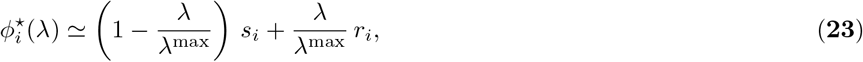

with 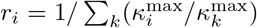, which is found by re-arranging terms in [**20, 21**]. Hence, the expression of a given enzyme is inversely proportional to its square root specificity at low growth and to its catalytic rate 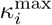 at high growth. Defining the ratio 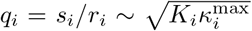, metabolic growth response distinguishes two classes of enzymes: rate-optimized enzymes (*q*_*i*_ *>* 1) are up-regulated, specificity-optimized enzymes (*q*_*i*_ *<* 1) are down-regulated upon nutrient limitation (Fig. 2, Fig. 4A). In networks with a sizeable proteome fraction 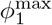, there is a small but systematic shift of the growth reponse pattern (SI).

The corresponding global resource allocation pattern is shown in Fig. 4C: a depletion of metabolites comes together with an expression shift from specificity-optimized to rate-optimized enzymes. These analytical predictions capture specific networks with physiological reaction parameters, not just ensemble averages (Fig. S4).

### Growth-dependent patterns in genome-wide protein expression data

A comprehensive test of our model requires system-wide kinetic data, 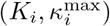, of metabolic reactions, which are currently not available. However, statistical evidence can be obtained from whole-genome proteomics data. Here we use a recent dataset of protein expression for 1743 genes in *E. coli* growing under different levels of glucose uptake limitation (24). For each gene, we plot the expression level at high nutrient supply, 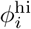, against the relative change upon nutrient limitation, 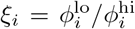 (Fig. 5). We find a negative correlation, 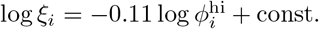, that is statistically significant (*p <* 0.03 under a null distribution with scrambled growth rates). This type of correlation can be explained by our model: high values of 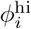 are correlated with low values of 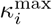, signalling predominantly specificity-optimized proteins (*q*_*i*_ *<* 1) that are down-regulated upon nutrient limitation (assuming no overriding correlation between *κ*^max^ and *K*). The signal identified in the proteomics data is likely to be diluted by several factors of network heterogeneity not included in the minimal model. Typical growth responses under nutrient limitation affect predominantly specific functional sectors and exclude others, such as metabolically inactive proteins (24, 42) (see also below). Specific functional groups involved in a common reaction, such as the ribosomal genes, are consistent with the model on average but do not show a within-group anti-correlation of ξ and 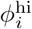 (Fig. 5). Other groups, including the pathways of tetrahydrofolate synthesis or glycolysis, involve reaction chains and show the predicted anti-correlation with pathway-specific offsets.

**Figure 5:**
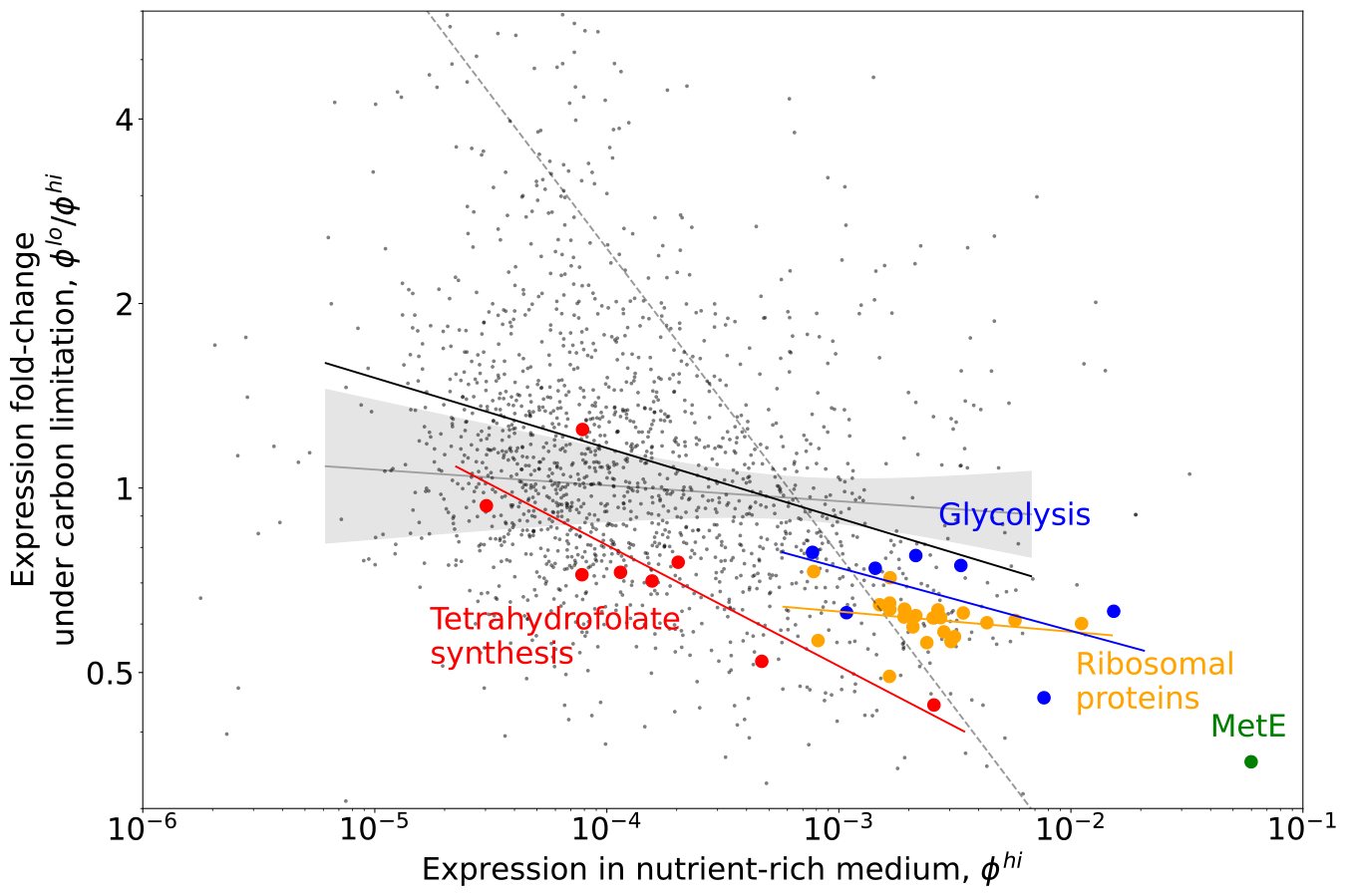
Protein expression changes under nutrient limitation. Expression fold change upon glucose uptake limitation, 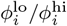 plotted against expression in a nutrient-rich environment, 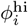, for 1743 proteins of *E. coli*; data from (24). These data show a negative correlation (solid black line) that is statistically significant (*p* < 0.03; the shaded region shows the range of slopes expected from data with randomized growth rates). The biochemical network model explains this correlation by the growth-dependent expression shift from specificity-optimized to rate-optimized enzymes (see text). Individual single-reaction pathways are consistent with the model in their average expression (orange: ribosomal proteins), multi-step pathways (blue: glycolysis, red: tetrahydrofolate synthesis (43)) show intra-group anti-correlations with pathway-dependent offsets.

### Growth response of functional sectors

How does the pattern of growth response generalize to more complex metabolic networks⋆ Here we consider a network with two uptake pathways, C and N, processing distinct nutrients both necessary for growth; their outputs feed into a single downstream pathway, A (Fig. 4D-F). This is one of the simplest networks with differentiation into functional sectors. In previous work, such sectors have been identified as coherent units of proteome resource allocation (44, 45). We study the three-sector network under limitation of nutrient C, assuming again an expression constraint 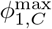 of the cognate uptake machinery. To obtain the growth-optimal resource allocation pattern shown in Fig. 4D-F, we employ Eq. [**6**] in each pathway, and numerically solve for optimal resource allocation at their junction. Sectors C and A show a common growth response similar to the single-chain system [**22, 23**]: metabolites are depleted, proteins experience an expression shift from specificity-optimized to rate-optimized enzymes. In sector N, metabolites are not depleted and enzymes are uniformly down-regulated, inducing a net resource shift from N to C and A (Fig. 4F).

Together, individual metabolite and enzyme levels depend on their functional sector and on the biochemistry of the local reaction step. Thus, our model produces growth response patterns with coherent shifts between sectors, as well as heterogeneous changes within sectors. This heterogeneity is consistent with available experimental data: multiple groups of proteins characterized by distinct regulatory responses to external perturbations are enriched in the same functional Gene Ontology-terms (24). Local biochemistry can also contribute to coherent shifts if kinetic parameters are broadly correlated with sectors, as suggested by recent empirical observations (46). A specific case in point are ribosomes, which have a small catalytic rate per unit of mass, 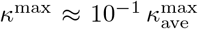 (1) and a small binding constant *K <* 10^*-*2^*K*_ave_ (47, 48), generating a sub-average growth response parameter *q <* 1 (SI). Hence, by local biochemistry, ribosomes are highly expressed in nutrient-rich media and down-regulated with decreasing growth; a similar example may be methionine synthase (MetE) (Fig. 5). Such low-*q* anabolic enzymes with large proteome fractions may contribute to the generic down-regulation of the A sector experimentally observed under multiple growth-limiting conditions (24, 42).

### Metabolic response by regulation

Can a metabolic system come close to an optimal growth state by local regulation of its enzymes (49)⋆ Specifically, we consider the response of a metabolic chain to nutrient limitation. This process requires growth-dependent shifts in metabolite and enzyme levels, 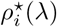 and 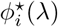, as given by equations [**22**, **23**]. Protein concentrations change by synthesis with rate *b*_*i*_ and by dilution, *ϕ*_*i*_ = *b*_*i*_ - λ*ϕ*_*i*_. In the simplest case of transcriptional regulation, the stationary level

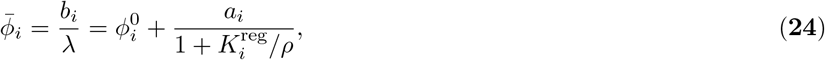

depends on the equilibrium binding of a transcription factor present at density *ρ* (SI and Fig. 6A). This regulation function has three parameters: the basal level 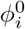, the regulatory amplitude *a*_*i*_, and the regulatory binding constant 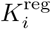. Here we assume a simple logic: at each metabolic step, the substrate regulates the transcription of its cognate enzyme (Fig. 6A). In this network, growth optimization by regulation depends on a remarkable property: with regulatory parameters depending only on local biochemistry,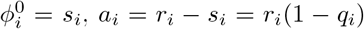, and 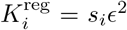, the stationary expression level [**24**] follows the optimal growth response, 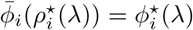, up to terms of order ϵ. This identity follows by inspection of equations [**22** – **24**]; see SI for details. As required for optimal growth response, rate-optimized enzymes (*q*_*i*_ *>* 1) are repressed, specificity-optimized enzymes (*q*_*i*_ *<* 1) are activated by their cognate metabolite. The adaptive dynamics of the metabolic state, S(*t*) = (*ρ, ϕ*)(*t*), consists of regulated changes of protein levels and correlated changes of metabolite levels given by mass conservation, equation [**3**]. Here we use metabolite-mediated regulation specified by the set of parameters ℛ_*m*_ = (ϕ^0^, *a, K*^reg^) for the entire network. Fig. 6B shows these dynamics under time-dependent nutrient limitation, *ρ*_1_(*t*), switching between subsequent stationary levels. Rate-optimized and specificity-optimized enzymes change in opposite directions and settle to a stable stationary state 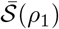 with growth rate 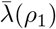 that matches the growth-optimal state in very good approximation,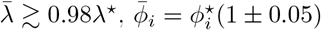,and 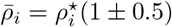 (Fig. 6C).

**Figure 6:**
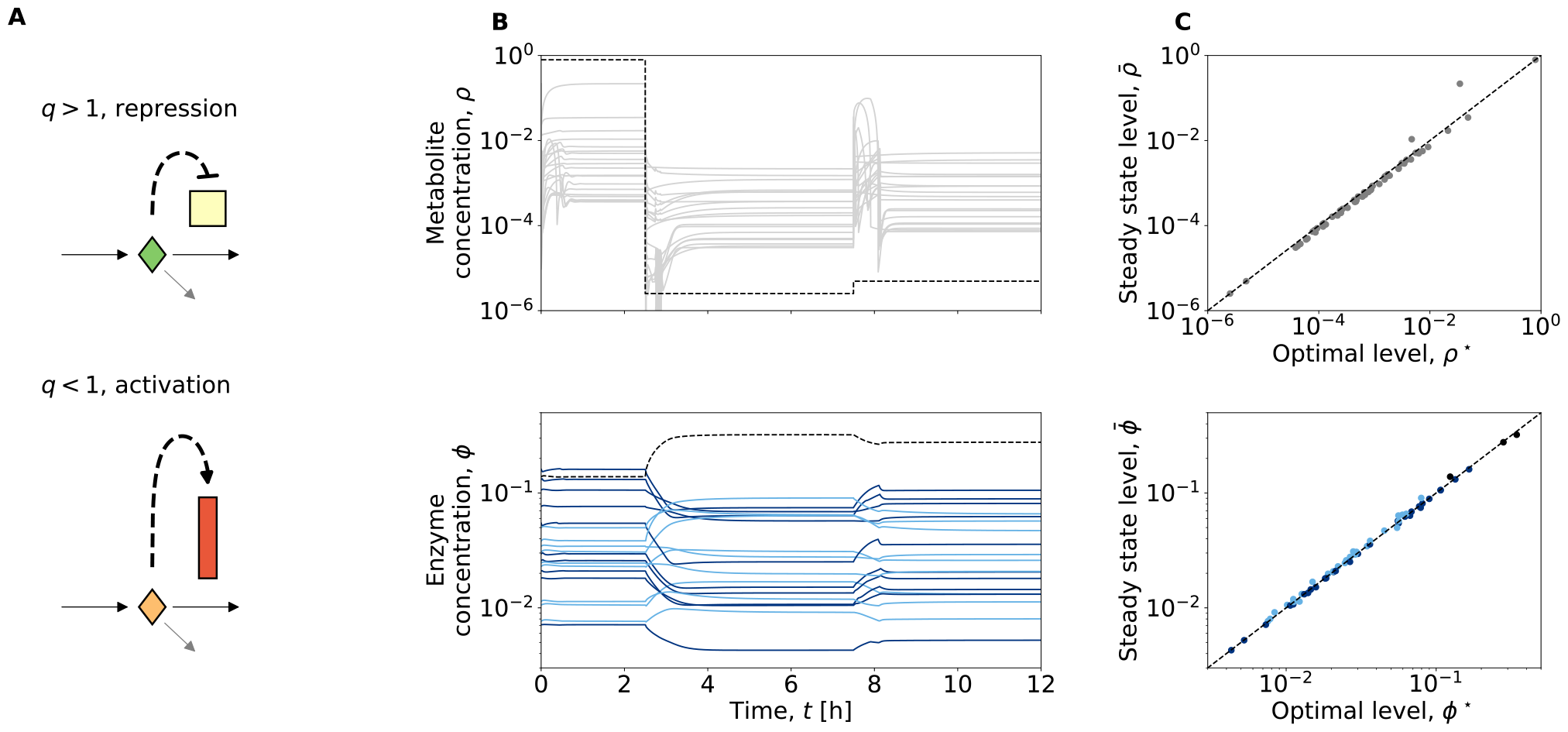
Metabolite-mediated regulation. (*A*) Logic of regulation: metabolites act as transcription factors for their cognate enzymes, inducing repression for rate-optimized enzymes (*q*_*i*_ > 1) and activation for specificity-optimized enzymes (*q*_*i*_ < 1). (*B*) Time-dependent metabolite levels, *ρ*_*i*_(*t*) (gray) nutrient concentration (dashed), and regulated protein concentrations ϕ_*i*_(t) (dark blue: specificity-optimized enzymes, q < 1; light blue: rate-optimized enzymes, q > 1, dashed: uptake enzyme) under time-dependent nutrient limitation, *ρ*_1_(t) (top, dashed). (*C*) Regulated late-time stationary levels, 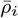 and 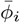, plotted against growth-optimal levels, 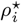 and 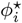.

More broadly, our analysis suggests that metabolite-induced regulation of enzymes can generate partial optimization of metabolic states under diverse conditions. In more complex networks, local regulation of each enzyme is not a plausible scenario. However, few, broad-acting transcription factors can generate coarse-grained optimization of the average expression for co-regulated target genes in pathways or functional sectors.

### Network evolution

On large time scales, metabolic networks change by protein evolution. Molecular changes affect local reaction parameters 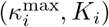, inducing changes in pathway usage and, ultimately, in network topology. Consider an enzyme mutation changing the affinity to its cognate metabolite, *K*_*i*_ *→K*_*i*_(1 + *δK*_*i*_). Its effect on growth is partially buffered by the physiological response of the network, which regulates the protein expression *ϕ*_*i*_ and changes the metabolite density *ρ*_*i*_. Our model predicts the strength of selection on network-buffered binding affinity changes,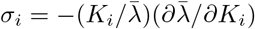, where we assume a regulation function ℛ reproducing near-optimal stationary states, 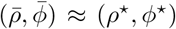, in the wild-type system (*κ*^max^,*K*). We obtain buffered selection coefficients,

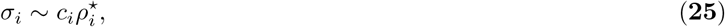

where *c*_*i*_ are growth-dependent factors of order 1 (SI). Two important consequences follow. First, given 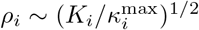, selection acts on enzyme specificities. In other words, the selection pressure to increase the affinity 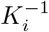 is higher for enzyme-metabolite pairs with low catalytic rate *κ*^max^, predicting a negative correlation between affinity and rate in evolved networks. Second, we can estimate the order of magnitude of selection by evaluating *ρ*_*i*_, as given by equation [**22**] at intermediate growth rates. In the mean-field approximation introduced above, we find *σ*_*i*_ *∼ϵ*^2^/*𝓁 ∼*10^*-*4^ at physiological network parameters *K*_ave_ ∼10^*-*4^ and *𝓁∼*20. Importantly, the buffered *σ*_*i*_ is at least two orders of magnitude lower than the growth effect of an affinity change for a single Michaelis-Menten pair, *σ*_0_ = (*K*/*J*)(*∂J*/*∂K*) = (1 + *ρ*/*K*)^*-*1^ at constant *ρ* [**1**].

The evolution of enzyme-metabolite binding is a mutation-selection-drift process (50) on a biophysical fitness land-scape (51–58), here of the form *f*(Δ*G*_*i*_) ∼ −*C*_*i*_ exp[Δ*G*_*i*_/2]), where Δ*G*_*i*_ = log(*K*_*i*_/*K*_0_) is the binding free energy and *C*_*i*_ is a constant. In such landscapes, mutation-selection equilibrium emerges at typical selection coefficients *σ*_*i*_ ∼ 1/*N*_*e*_, where *N*_*e*_ is the effective population size (58–60). Hence, the strength of selection on *K*_*i*_ is consistent with evolutionary equilibrium at *N*_*e*_ ∼ 10, in tune with the effective population sizes obtained for microbial systems evolving under clonal interference of protein phenotypes (in such systems, 1/2*N*_*e*_ is the average number of generations between subsequent selective sweeps) (58). In turn, this implies that physiological values of the metabolome-proteome mass ratio, *M*_*R*_/*M*_*P*_, are set by an evolutionary equilibrium where the selective cost of metabolite dilution balances with mutations degrading enzyme-metabolite binding.

## Discussion

Here we have developed a model for resource allocation in metabolic networks that integrates the kinetics of local reaction steps, the global structure of the network, and external nutrient constraints to predict enzyme and metabolite levels in growth-optimal network states (Fig. 2). Protein expression levels are set predominantly by local biochemistry: levels at high growth rates are set by catalytic rates, 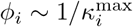 the response to growth limitation is linear with a slope depending on the response parameter 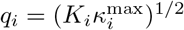. This pattern is in broad agreement with empirical observations in metabolic networks. First, the model explains the heterogeneous growth response of enzymes with similar functions (24). Second, it predicts finite protein expression levels in the limit of low growth rates, specifically for the ribosome; different mechanisms for this residual expression have been discussed in the recent literature (34, 42, 61, 62). Third, the linear growth response of protein levels derived here reproduces the linear pattern found in previous work at the level of functional protein sectors (15, 44, 62). However, the coherent growth response of broad sectors (related, e.g., to limitation of a particular nutrient influx), is not a generic outcome for growth-optimized networks. Our analysis suggests two possible mechanisms generating sector responses: broad correlations of biochemical parameters and network position (46) or deviations from fine-grained optimality by pleiotropic, rather than gene-specific, regulation.

Metabolites are predicted to show a nonlinear, position-dependent depletion of steady-state levels in response to growth limitation. Moreover, we show that metabolite levels depend on a local cost parameter *β*_*i*_ that summarizes constraints affecting the upstream metabolic chain (here, nutrient limitation). Such growth laws for metabolites have not yet been tested systematically by experiment; available data suggest that some but not all metabolites are depleted by growth limitation (25, 63, 64). Deviations from model predictions may signal toxicity or enhanced efflux of specific metabolites. They may also reflect an increase in *β*_*i*_ caused by bottlenecks in upstream reactions or nutrient conditions, in tune with an observed decrease in metabolite concentrations on poor carbon sources (16). More broadly, our model can serve as a starting point to describe the quantitative organization of the metabolome, which has remained elusive so far (39).

A mechanistic model of metabolic networks including enzymes and metabolites is also a prerequisite for predicting network dynamics by regulation and, on longer time scales, by evolution of reaction rates and binding constants. Here we have shown that a simple regulatory scheme based on feed-forward regulation – each metabolite regulates the enzyme of the next reaction step – can closely approximate optimal resource allocation under nutrient limitation (Fig. 6). Thus, our model highlights the role of metabolites as regulatory signals and paves the way to design regulated synthetic networks that maintain optimality under variable input (49, 65).

Evolution acts on network states buffered by regulated physiological response. The resulting selection coefficients for local changes in network biochemistry, which become computable from our model, are lower than in the absence of network buffering. Specifically, we have shown that metabolite dilution induces system-wide selection pressure on enzyme-metabolite binding. Individual enzymes evolve on approximately independent fitness landscapes depending on their target affinity. Evolutionary equilibria on these landscapes can explain the order of magnitude of physiological enzyme-target affinities, which are well below the biophysical bound set by substrate diffusion (37), as well as the global mass ratio of metabolome and proteome. In contrast, interventions like antibiotics can impair specific enzymes in a stronger way, often beyond the limits of regulatory buffering (66), generating stronger selection pressure on evolutionary network changes (18, 19). Understanding this interplay of regulation and evolution is crucial for the design of efficient antibiotic interventions into microbial metabolism.

## Acknowledgements

We thank Tobias Bollenbach, Laura Collesano, Martin Lercher, and all members of the Bollenbach and Lässig groups for helpful discussions. This work was supported by Deutsche Forschungsgemeinschaft Grant SFB 1310 (to ML). FP acknowledges funding by Human Technopole.

## Supporting Information

### Maximization of balanced growth rate by resource allocation

#### Scaling of variables

Bacteria reallocate metabolic resources under environmental challenges. We explore growth-optimal resource allocation in a structured network of biochemically heterogeneous enzymes (Fig. 1) in variable environments. In contrast to previous proteome sector models (44), our model concurrently accounts for both metabolites and enzymes. As a result, our model introduces second-order rate constants that describe reactions with multiple reaction partners, such as enzyme-substrate interactions. The rates and concentrations in our model exhibit a different scaling with the reaction volume when enzyme and substrate amounts are held constant. To include this scale in our model, we introduce the parameter

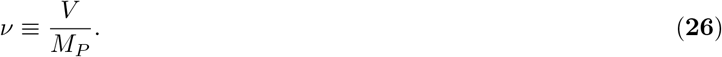

Here, *V* is the total intracellular volume, and *M*_*P*_ is the total mass of protein. Empirical observations suggest that this fraction remains approximately constant throughout the cell cycle and across different environments (15). The parameter ν enables converting mass concentrations *m*/*V* into dimensionless mass fractions *m*/*M*_*P*_. A metabolite mass fraction *ρ* is defined as *ρ* = *M*_*S*_*νc*, with the classical concentration *c*, in units of mol L^*-*1^, and the molar mass of the metabolite *M*_*S*_. For simplicity, we apply identical unit conversions to the extracellular nutrient concentration *ρ*_1_. Similarly, a protein mass fraction *ϕ* corresponds to *ϕ* = *M*_*E*_*νc*_*E*_ with the classical concentration *c*_*E*_ (in units of mol L^*-*1^) and the molar mass of the protein *M*_*E*_. In this unit system protein mass fractions in a cell of *ℓ* proteins must sum up to 1,

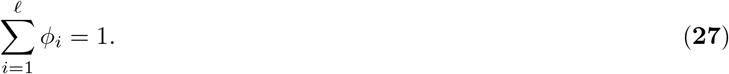

Flux units are accordingly chosen to directly express the time derivatives of mass fractions as flux summations, *dρ/dt* = *j*_production_ − *j*_consumption_ − *j*_dilution_. The conversion also extends to kinetic rate constants. For instance, a Michaelis-Menten reaction depicts an enzyme with a mass fraction *ϕ* consuming a metabolite *ρ* at a rate *j* = *κ*^max^*ϕ*/(1 +*K*/*ρ*). Here *K* is a dimensionless Michaelis constant, linked to the classical Michaelis constant 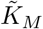 (in units of mol L^*-*1^) via 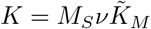. *κ*^max^ is the enzyme’s turnover number in this unit system and relates to the classical turnover number *k*_cat_ via *κ*^max^ = (*M*_*S*_/*M*_*E*_) ·*k*_cat_. This unit conversion allows us to write a compact and (apart from time) dimensionless derivation in the following. In this unit system, the growth rate is compactly represented as the mass-normalized flux through the protein biosynthesis reaction, which we take as the last reaction of the metabolic cascade, *j*_*𝓁*_,

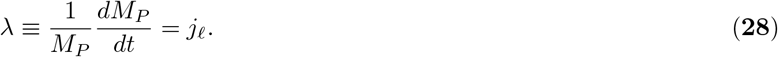

Furthermore, we define the following extensive variables to simplify notation throughout: *R*_*i*_ ≡ *ρ*_*i*_*M*_*P*_, *M*_*i*_ ≡ *ϕ*_*i*_*M*_*P*_ and *J*_*i*_ ≡ *j*_*i*_*M*_*P*_.

#### Independent variables: metabolite fluxes and nutrient concentration

We restrict our analysis to balanced growth states. This is exponentially growing, metabolic steady-states that obey metabolite mass conservation in each metabolic reaction. In balanced exponential growth (averaging over cell-cycle dependent and stochastic variation), no intracellular metabolite concentration may change over time (Eq. [**3**] must be zero for *ρ*_*k>*2_). Furthermore, all 𝓁 reactions must fulfill the kinetic rate law [**1**]. As a result, only a specific subset of all possible metabolic states, denoted by 𝒮, is viable during balanced growth. We demonstrate that, given a chain of enzymes with kinetic parameters *κ*^max^ and *K*, any metabolic state during balanced growth is fully determined by its metabolite fluxes *J*, and a nutrient environment *ρ*_1_: inserting Eq. [**2**] into the balanced growth condition (Eq. [**3**] being zero) and solving for *ρ*_*i*_ yields all but the first metabolite concentration as a function of the fluxes (and vice versa [**4**]),

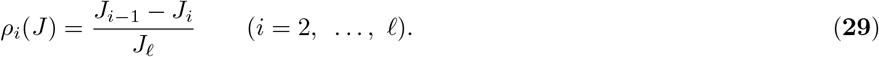

The environmental nutrient concentration *ρ*_1_ is not constrained by balanced growth but by the experimental setting.

We next compute each individual protein mass according to the kinetic rate laws,

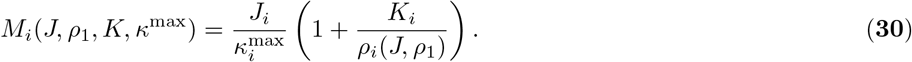

By summation, the total protein mass can easily be computed, and subsequently *R* from *ρ, ϕ*_*i*_ from *M*_*i*_, *j* from *J*, as well as the growth rate of the metabolic state in balanced growth. Taken together, growth rate can be expressed as a function of all metabolic fluxes, the nutrient environment, and the kinetic constants.

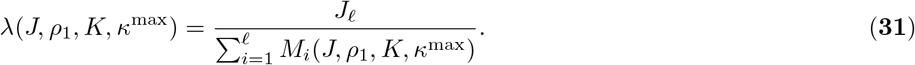

#### Optimality of metabolic states

Akin to flux balance analysis (14), we employ constraints to identify physiological solutions. A key constraint is the constant intracellular protein density ν^*-*1^, which is well established empirically (1, 15, 26–28). Due to this constraint, we can identify a metabolic state 𝒮^⋆^ that optimizes growth rate *λ*: if an enzyme’s concentration is too low, it limits growth through a metabolic bottleneck. If its concentration is too high, it demands a reduction of other enzyme’s concentrations, which also impairs growth.

We focus only on the regulatory problem of finding optimal metabolic states 𝒮^⋆^ for a given set of kinetic constants (*κ*^max^, *K*). We assume that kinetic constants evolve on a slower timescale, in contrast to simultaneous evolutionary optimization of kinetic constants and expression levels. To shorten the notation, we drop the kinetic constants from all arguments below. Optimal balance growth states are defined as

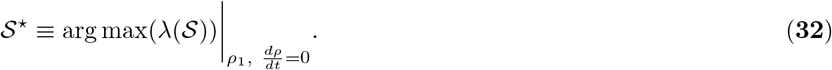

As shown in the preceding paragraph, metabolic states that fulfill balanced growth and the kinetic rate laws are fully determined by their fluxes *J* and a nutrient environment *ρ*_1_ [**31**]. We thus rewrite the previous equation,

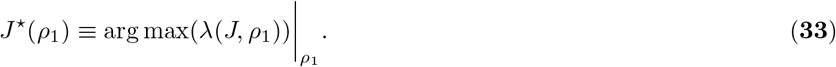

By construction, this optimal state can grow exponentially with time-independent metabolite concentrations. For any system size *M*_*P*_, at least one optimal state exists. We here set the system size implicitly by choosing a certain *J*_*𝓁*_. Maximal growth rate is achieved if the protein mass necessary to produce this biomass production flux *J*_*𝓁*_ is minimal; see Eq. [**2**]. Thus, we can simplify the optimality condition further and obtain Eq. [**5**] - optimality is achieved only if

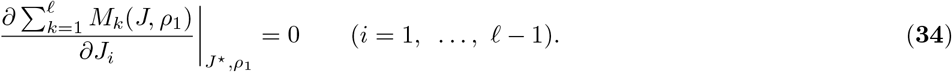

This principle resembles Enzyme Cost Minimization (36), which has been used to numerically find growth rate maximizing resource allocation in Flux-Balance-Analysis when considering reversible kinetic rate laws but no dilution of metabolites. Here we instead focus on growth maximization in a model that explicitly takes metabolite dilution into account, but no reversibility. We proceed analytically. A restricted analysis of this kind was previously pursued by Ehrenberg and Kurland (30). Beyond their analysis, we quantify the relative influences of enzyme biochemistry and network properties on resource allocation to individual enzymes and metabolites and analyze how these influences change under environmental challenges.

### Iterative construction of the optimal metabolic states in longer chains

#### Optimal resource allocation in coarse-grained metabolic models

A cell model consisting of just two proteome sectors (𝓁 = 2) - with catabolic enzymes *M*_1_ and anabolic enzymes *M*_2_ - has been employed successfully to infer proteome resource reallocation in response to changes in the nutrient environment *ρ*_1_ and the catalytic rate of the anabolic sector 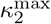 (15). For such a short chain, the optimality condition [**33**] has also been solved analytically, including both the proteome and the metabolome (33).

#### Producing the first substrate of the chain from a new environmental precursor

Longer chains feature many non-linearities and feedback loops, and can not easily be solved analytically anymore. To find the optimal metabolic states 𝒮 ^⋆^ (*J* _*𝓁*_, *ρ*_1_) for longer chains, we identify an iterative scheme for constructing optimal metabolic states of a chain with *𝓁* +1 enzymes from the optimal metabolic states of a chain with *𝓁* enzymes. Suppose the optimal metabolic state 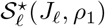 is known for a given chain defined by its 2 *𝓁 kinetic* constants in an environment *ρ*_1_. We extend the given chain by a single enzyme with its kinetic constants, a protein mass 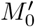, catalyzed flux 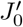 and a new environmental precursor concentration 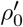 instead of *ρ*_1_. The new enzyme produces the first substrate of the shorter chain (Fig. S1). We indicate variables of the extended chain by a dash ^*‘*^, and drop the superscript ^⋆^ to simplify the notation in this section. Below, all state variables denote their growth-optimal values.

#### Recursion relations for state variables

An extension that adds one additional flux 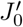 but conserves all fluxes of the shorter chain 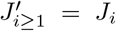, also conserves all downstream concentrations 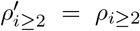 and thus all downstream protein masses 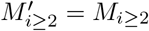, according to Eqs. [**29**] and [**30**]. An extension of the chain by a single enzyme with arbitrary flux therefore only changes *ρ*_1_ to a new 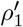 for which mass conservation now applies, correspondingly *M*_1_ to a new 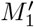, and adds 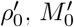 and 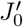.

These five new or modified state variables are linked by five equations: two kinetic rate laws (Eq. [**1**] for *i* = 0 and *i* = 1), a mass conservation constraint (Eq. [**29**] for *i* = 1), and two conditions on optimality (Eq. [**34**] for *k* = 0 and *k* = 1). These five equations can be solved for the five state variables. The solution is jointly given by

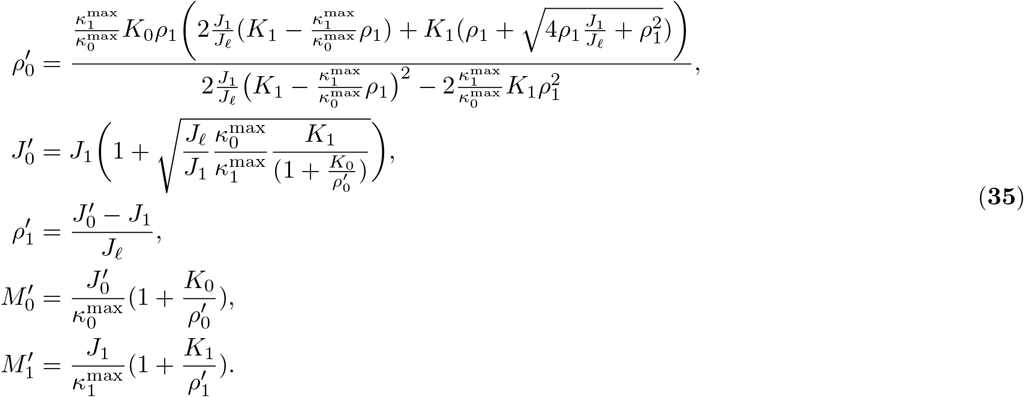

A single step of this iteration is illustrated in Fig. S1. All other conditions on optimality (Eq. [**34**] for *k ≥ 2*) are fulfilled by the extended chain if they were fulfilled by the shorter chain because the protein masses 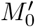 and 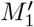 do not depend on down-stream fluxes 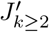 in a system of irreversible enzymes 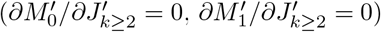, while the required protein mass downstream 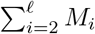 only depends on downstream fluxes in a manner independent of the extension 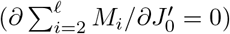. Precisely this property of the model cell allows for a local construction of a new chain start that again yields an optimal chain, rather than requiring a full adjustment of the whole chain.

#### Direct iteration of fluxes

Computationally, it is practical to calculate the concentration of an external nutrient just once. Instead of iteratively determining the five modified state variables in each step, fluxes alone can be iterated according to the optimality criterion of Eq. [**34**]. To derive an explicit formula, we consider a variation of a single flux *J*_*i*_ + *δJ*_*i*_ (with i = 2, …, *𝓁* − 1), while keeping all other fluxes constant, as required by the optimality criterion. Among all mass conservation criteria [**29**], only those for *ρ*_*i*_ and *ρ*_*i*+1_ depend on *J*_*i*_. We find *ρ*_*i*_ + *δρ*_*i*_ = (*J*_*i-1*_ *-* (*J*_*i*_ + *δJ*_*i*_))/*J*_*𝓁*_ and *ρ*_*i*+1_ + *δρ*_*i*+1_ = (*J*_*i*_ + *δJ*_*i*_ *J*_*i*+1_)/*J*. Together, these changes in metabolite concentrations and the change in flux affect enzyme masses according to the Michaelis-Menten relation [**1**]. Inserting the changes derived above yields mass changes under variation of a single flux around the optimal state,

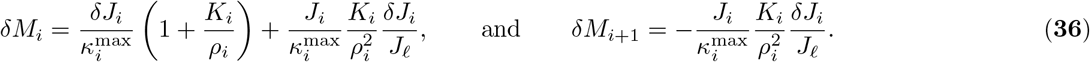

To summarize, for *i ≥* 2, only *ρ*_*i*_ and *ρ*_*i*+1_ (see Eq. [**29**]) depend on *J*_*i*_. Only *M*_*i*_ and *M*_*i*+1_ depend on either *J*_*i*_, *ρ*_*i*_ or *ρ*_*i*+1_. The respective changes are illustrated in Fig. S1C. The optimality criterion [**34**] can now be drastically simplified to 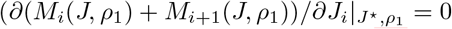 for *i* = 2, …, *𝓁*− *1. By* inserting the derivatives [**36**], and expressing metabolite concentrations through fluxes [**29**], we find Eq. [**6**], which can be solved for *J*_*i-1*_,

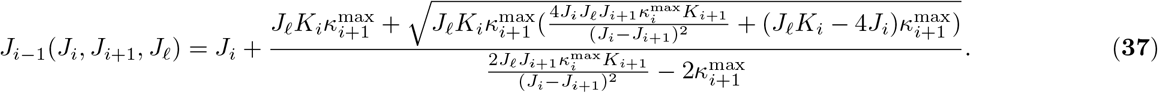

The result only depends on downstream fluxes *J*_*q≥i*_ and kinetic constants. From a starting condition *J*_*𝓁*_ and *J*_*𝓁-*1_ all fluxes in the chain can be determined iteratively. Only in the last iteration step, a matching nutrient concentration is computed. It is given analogously to Eq. [**35**] by

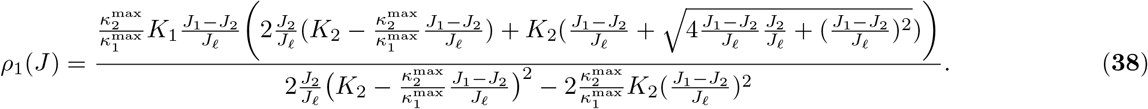

At this point, Eqs. [**29**] and [**30**] can again be used to compute all other metabolic state variables of 𝒮^⋆^.

#### Matching the environmental nutrient concentration

Starting from a minimal chain that consists of only two enzymes, and all its possible optimal metabolic states 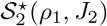 we can construct optimal metabolic states of a three-enzyme-chain. By shifting indices, we can iteratively determine optimal metabolic states 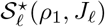 for any longer chain.

In many biologically relevant scenarios, a boundary condition to be fulfilled is the environmental precursor concentration *ρ*_1_. However, we here construct a suitable environmental nutrient concentration *ρ*_1_ iteratively from the environmental nutrient concentration of the initial two-enzyme chain. It can easily be seen from Eq. [**35**] that if any positive *ρ*_1_ was possible for the shorter chain, any positive 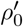 can be constructed for the elongated chain. Thus, the iterative method we present here can construct an optimal resource allocation for any environmental nutrient concentration. Unfortunately, it is not clear from the outset of the iteration, which environmental precursor concentration of the two-enzyme chain yields the environmental precursor concentration for the *𝓁*-enzyme-chain set by the biological scenario of interest. We solve this question numerically, whenever a defined nutrient environment is required.

#### Uniqueness of optimal resource allocation

Any local extremum of growth rate found in this manner is a global optimum because the solution of the chain of two enzymes contains all optima of the chain of two enzymes. In each step, only one physically meaningful solution (with all metabolic state variables larger than zero) of the set of five equations solved for the extension of the chain exists. Thus, the optimal states *J*^⋆^ determined here are indeed the only sets of fluxes that fulfill Eq. [**5**]. No physiological growth maxima exist at extremal values of any *J*_*i*_. Consequently, the optimal metabolic state 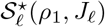 is unique and all optimal metabolic states of linear pathways can be constructed by the iterative method.

### Local properties of optimal resource allocation

#### Metabolite costs increase along the metabolic chain

We next show how optimal resource allocation in a network relates to the individual enzyme-substrate pair. To this end, we switch from a variation of a single flux *J*_*i*_, which we used to construct growth-optimal balanced growth states above, to a variation of a single metabolite concentration *ρ*_*i*_ + *δρ*_*i*_. We again derive the mass changes under such a variation. Eq. [**4**] connects metabolite concentrations and fluxes in balanced growth states. It indicates that a variation of a single flux can be constructed if all fluxes upstream of reaction *i are incr*eased by a small *δJ*. We insert all changes in fluxes into the Michaelis-Menten relationship to obtain changes in enzyme masses,

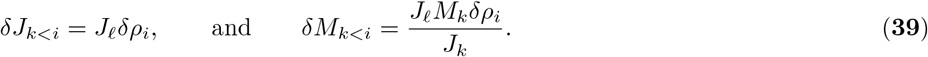

This variation is illustrated in Fig. S1D. Here we implicitly assumed that the environmental nutrient concentration *ρ*_1_ does not depend on the uptake flux *J*_1_. This may in general not be true under nutrient limitation - for example in a chemostat setting, in which increased nutrient uptake reduces the steady-state nutrient concentration - and should only be understood as a limiting case in the nutrient-rich regime. We discuss the effect of nutrient limitation in more detail below. The focal reaction is also affected by the variation in *ρ*_*i*_ - not by a change in flux, but by the change in substrate concentration. We will investigate *δM*_*i*_ in more detail below. For a constant protein production flux *J*_*𝓁*_ and nutrient environment *ρ*_1_, the total protein mass in the cell must also be minimal under variation of a single metabolite concentration (that is under a linear combination of changes in fluxes [**39**]). We simplify based on the changes in masses derived above to obtain Eq. [**8**],

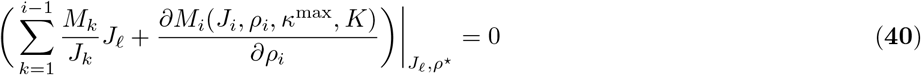

In the optimal state, 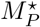 must be a constant to first-order approximation in any of the variations discussed here. We can therefore equivalently write

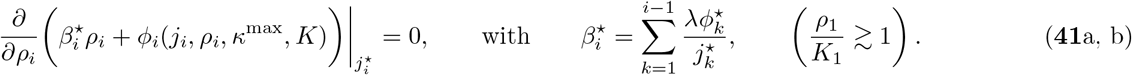

Eq. [**41**a] corresponds to Eq. [**9**]. The constant [**41**b] (which we define explicitly as independent of the variation in *ρ*_*i*_) corresponds to the second term of Eq. [**10**]. The full form of Eq. [**10**] will be derived below. We obtain a result similar to the minimization of the joint mass concentration of an enzyme and its substrate at constant flux proposed by Dourado et al. (16), 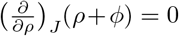, which explains experimentally observed relations between enzyme and substrate mass concentrations. In contrast to Dourado et al. we here minimize the total *protein* mass at a fixed *protein* production flux. Our optimization directly follows from growth-optimality: In balanced exponential growth, each constituent of the cell must double at the same rate. Applied to protein, this means that growth rate is given by the protein production flux *J* divided by the total protein mass *M*_*P*_ of the cell. At fixed *J* _*𝓁*_, *M*_*P*_ must be minimal to achieve optimality. From the outset, it is not clear how such a minimization of the protein mass relates to the metabolite level. We here show that a joint minimization of protein and weighted metabolite mass follows from the optimization of growth rate. The weight 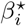 must be derived for a given metabolic network, and is generally different from 1. We retrace the derivation of the scaling law discovered by Dourado et al. (16) with the additional cost 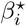. Eq. [**30**] can be rewritten intensively as 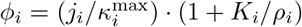, and inserted into Eq. [**41**a]. Evaluating the derivative and rearranging yields Eq. [**12**], and by re-inserting into Eq. [**30**], we find Eq. [**11**].

### Physiological model parameters

#### Growth costs of metabolites and dilution strength *ϵ*

Eqs. [**11**] and [**12**] must ensure that all proteome mass fractions sum to one. This property can be used to establish a parameter scale, which distinguishes between physiological parameterizations that approximately obey a production-consumption balance in each reaction and non-physiological parameterizations that are dominated by dilution. We insert Eq. [**11**] into [**12**] and solve for 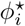,

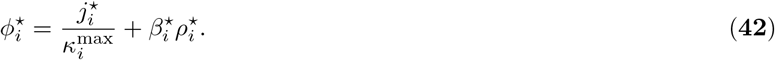

Summing over all proteome mass fractions [**27**], and inserting mass conservation [**4**], we derive

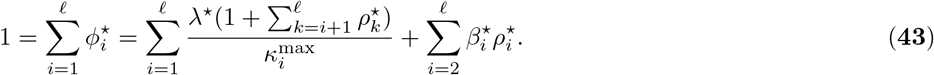

We insert the definition of metabolite costs [**41**b] (again in the nutrient-rich regime) to derive 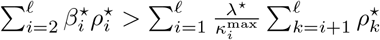, and establish two bounds on growth rate,

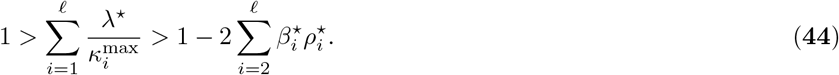

Growth rate must be smaller than the inverse of the sum of all inverse efficiencies, termed *λ*^*max*^, which would be the growth rate if all enzymes could work at maximal saturation without dilution losses. At the same time, scaled growth rate is reduced by less than 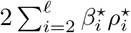. By employing again the definition of metabolite costs [**10**], and 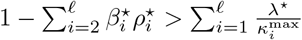 [**43**], we find that 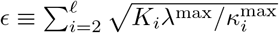 is an upper bound for 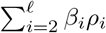. Below, we show that physiologically parameterized cells exhibit a small value of *ϵ* ≈ 10^−1.3^. Thus, their growth rate in nutrient-rich environments is predominantly determined by enzyme efficiencies.

#### Physiological kinetic parameters of enzymes

To estimate the value of *≈* we use available data for kinetic rate constants. Bar-Even et al. (37) found that enzyme kinetic parameters across numerous species tend to follow a log-normal distribution. Restricting their analysis to *E. coli* enzymes yields distributions of similar mean and variance (see tab. 1). We leverage their results to approximate *K* and *κ* ^max^ distributions for model parameterization. To convert units as described above, we estimate the average molecular weight of substrates *M*_*S*_ based on the molecular weight of glucose and amino acids (BNID: 104877); the volume per mass ratio *ν* [**26**] as the inverse of the protein mass per volume density in *E. coli* computed from the product of the dry mass protein fraction of *E. coli* (BNID: 101955) and its dry mass density (BNID: 109049); and the average protein mass *M*_*E*_ using the average protein length (BNID: 108986) and the average molecular weight of amino acids in *E. coli* (BNID: 104877). We disregarded any molar mass variability for substrates and enzymes on the grounds they are minor relative to the intrinsic variation of the kinetic parameters 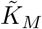 and *k*_*cat*_ and potentially correlated with one another and/or kinetic parameters.

#### Variability in *κ*^max^ and central metabolism dynamics

In comparison to a real cell, our model incorporates a reduced spectrum of enzyme types, namely those of a single pathway converting environmental nutrients to new protein biomass. In a real cell, enzymes unaccounted here convert metabolites through parallel, interconnected pathways and generate a variety of intermediary metabolites. In addition to a larger variety in protein precursors, these pathways also synthesize other cellular biomass components such as nucleic acids and lipids. Given the large spectrum of enzymes, the average proteome mass fraction of an individual enzyme type remains small. No singular enzyme mass fraction takes over the entire proteome, despite significant variability in catalytic rates.

To achieve physiological growth rates and prevent a singular enzyme from completely dominating mass allocation in our model, we use a distribution of enzyme efficiencies *κ*^max^ with lower mean and variance in comparison to the values found by Bar-Even et al. This choice addresses the reduced spectrum of enzyme types and is further motivated by evolutionary arguments: here we study central metabolism, which covers the conversion of environmental nutrients into new protein biomass. Central metabolism enzymes likely undergo more stringent selection than enzymes in secondary metabolism, resulting in decreased variance in their efficiencies. In contrast, we maintained the natural variance in substrate affinities and thereby maintained a near-physiological variance in enzyme qualities.

To estimate the chain length of the metabolic network in a living cell, we focus on the conversion of an environmental carbon source into new protein biomass. This pathway likely contributes most to the overall protein biomass production of the cell. Environmental carbon sources are converted into different amino acids via different pathways. To estimate the physiological pathway length, we use the production of glutamine from glucose, which consists of 17 enzymatic steps connecting the following metabolites:

Glucose → Glucose-6-phosphate → Fructose-6-phosphate → Fructose-1,6-biphosphate → GADP/DHAP → 1,3-Biphosphoglycerate → 3-Phosphoglycerate → 2-Phosphoglycerate → Phosphoenolpyruvate → Pyruvate → Oxalacetate/Acetyl-CoA → Citrate → Isocitrate → 2-Oxo-glutarate → Glutamate → Glutamine → Glutamineacyl-tRNA → Protein biomass

Other amino acid biosynthesis pathways include fewer (e.g. Serine), equally many (e.g. Threonine), or more enzymatic steps (e.g. Arginine). Alternative carbon sources may require a different number of enzymatic steps to produce pyruvate. Some enzymes may require additional modifications after protein synthesis. We here use *𝓁* ≈ 20 as an order of magnitude estimate for growth in amino-acid-free growth media.

#### Stochastic model parametrization

While kinetic properties of individual enzymes become increasingly available, only a minority has been measured to date. Additionally, enzymes with multiple substrates or otherwise complicated reaction kinetics may not exhibit a clearly defined substrate affinity. For these reasons, we do not parameterize our model based on available data for the specific enzymes. Instead, we parameterize our model stochastically, drawing both substrate affinities and kinetic constants independently from the log-normal distributions given in table 1. We thereby also disregard any bias in biochemical qualities along the metabolic cascade in our model. We note, however, that ribosomes exhibit a particularly high molecular mass but only an average catalytic constant *k*_*cat*_ *≈* 20 aa s ^− 1^ (1). Together with the average molecular weight of amino acids, these properties yield a distinctly low catalytic rate of *κ*^*max*^ = 10^0.9^ s^−1^. Furthermore, they operate very close to their maximal speed in the nutrient-rich regime (47, 48) despite typical aminoacyl-tRNA concentrations in the µM-concentration regime (BNID: 105275). We therefore estimate a strong ribosomal substrate affinity *K*^−1^ *> 10*^6^. Both properties are the sign of a specificity-optimized protein. As outlined further in the main text, we hypothesize that the ribosome is part of a coherently regulated group of specificity-optimized enzymes that lie downstream of most nutrient limitations.

**Table 1.** Distributions of enzyme kinetic parameters.

| | $\tilde{K}_M^*$ | $k_{\text{cat}}^*$ | $K = M_S \nu \tilde{K}_M$ | $\kappa^{\text{max}} = \frac{M_S}{M_E} k_{\text{cat}}$ |
| --- | --- | --- | --- | --- |
| All enzymes | $10^{2.0 \pm 1.2} \mu\text{M}$ | $10^{1.1 \pm 1.4} \text{s}^{-1}$ | - | - |
| *E. coli* | $10^{2.0 \pm 1.1} \mu\text{M}$ | $10^{1.1 \pm 1.3} \text{s}^{-1}$ | $10^{-4.2 \pm 1.1}$ | $10^{2.1 \pm 1.3} \text{h}^{-1}$ |
| Model cell | - | - | $10^{-4.2 \pm 1.1}$ | $10^{1.6 \pm 0.4} \text{h}^{-1} **$ |

To convert to the described unit system, we estimate $M_S \approx 10^2 \text{ g mol}^{-1}$ , $\nu \approx 6 \text{ mL g}^{-1}$ and $M_E \approx 3.6 \cdot 10^4 \text{ g mol}^{-1}$ .
\*: based on data from Bar-Even et al. (37).
\*\*: reduced mean and variance to account for the reduced number of enzymes in the model, see below.

#### Environmental nutrient concentrations

Amino-acid-free laboratory media for bacterial growth such as M9 typically contain between 0.*1g L*^−1^ and 1 g L^−1^ glucose. The glucose concentration 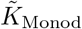 at which the growth rate of *E. coli* is half-maximal is on the order of 1 mg L^−1^ (BNID: 111048). We here assume - and later show that this assumption is in line with our model’s predictions - that the environmental nutrient concentration at which growth is half-maximal is on the order of the substrate affinity of the nutrient uptake enzyme, 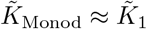. This suggests that nutrient uptake enzymes are typically highly saturated in nutrient-rich laboratory media. Based on these typical values, we specifically use *ρ*_1_ = 1000 *K*_1_ to describe nutrient-rich media. This value reflects all properties of the limit *ρ*_1_ well but is numerically more easily accessible than the limit itself.

### Global properties of optimal resource allocation

#### Physiological enzyme saturation

We compute the inactive proteome fraction *ϕ*_*inactive*_ (i.e. enzymes not currently engaged in catalysis due to a lack of substrates) as 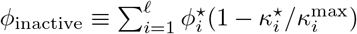. Inserting the definition of *κ* _*i*_ and Eq. [**12**], this yields

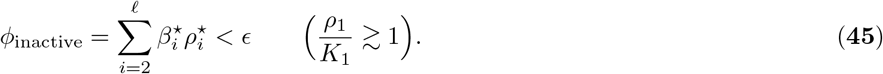

Enzymes of physiologically parameterized cells in nutrient-rich environments are typically highly saturated - on average to a level larger than 1 − *ϵ*. Our model thus explains solely from the physiological distribution of enzyme kinetic parameters, why enzymes are typically well-saturated in nutrient-rich environments (38).

#### The cellular metabolite mass fraction in the homogeneous chain

Consider a homogeneous chain (which consists of enzymes with equal kinetic constants *κ*^max^ and *K* only) in the nutrient-rich regime. In the physiological parameter regime, such a chain fulfills 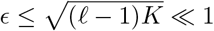. In the limit of small dilution strength *ϵ*, Eqs. [**1**], [**10**], [**11**], [**12**] and [**29**], are then jointly solved by

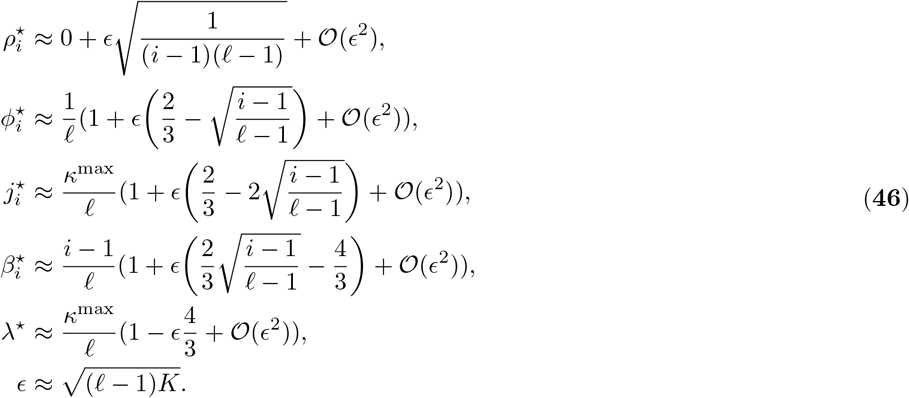

We here use 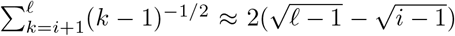 and 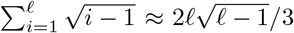, which are motivated by the corresponding integrals. With these approximations, Eqs. [**46**] form a consistent solution of microbial resource allocation to first order in *ϵ*, and the limit of vanishing *ϵ* constitutes the limit of vanishing dilution fluxes. In such a homogeneous chain, we can then compute the cellular mass fraction as *M*_*R*_/*M*_*P*_ = (*j*_1_ −*j*_*𝓁*_)/*j*_*𝓁*_ *≈*2*ϵ* [**18**], which is itself on the order of *ϵ*. At the same time, the growth effect of metabolite dilution is estimated [**46**] to be on the order of 4*ϵ*/3, which stems approximately equally from proteome resources required for unsaturated enzymes and proteome resources required to balance metabolite dilution fluxes [**44**].

#### Resource allocation in heterogeneous chains

To extend our analysis to heterogeneous chains, we replace position by the continuous variable *β*., and sums over position by integrals over 𝓁*dβ*. For a compact notation we define the relative specificity *sβ*.,

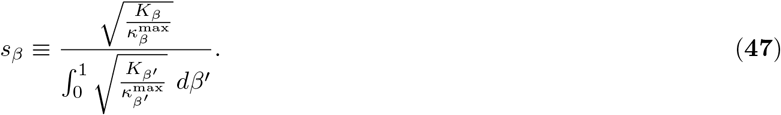

The central equations for *ρ, j* and *ϕ* then yield a self-consistent solution even in heterogeneous chains (under the additional assumptions that *β* is not correlated with other variables and fluctuations in other variables average out efficiently even across small intervals in *β* - i.e. any integral ∫_*L*_ *s*_*β*_. *dβ*. ≈ *L*):

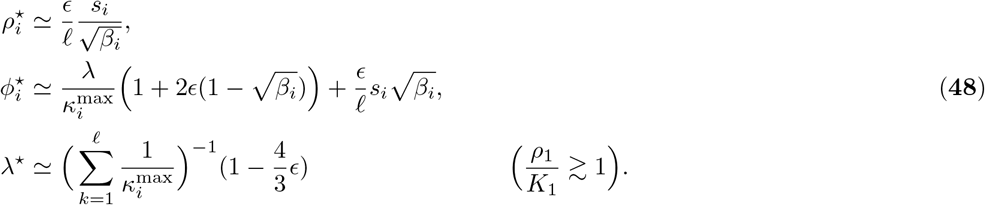

For sufficiently small variations of efficiencies and affinities around their respective mean, this derivation - together with the mean-field approximation for *β*._*i*_ [**46**] - provides a reasonable mean-field approximation of heterogeneous chains (Fig. S4). Systematic deviations introduced by physiological enzyme heterogeneity are small, as well as typical stochastic deviations. We note however, that this is not true for all heterogeneous parameterizations in the case of the cellular metabolite mass fraction. In heterogeneous chains, no upper limit on the metabolite mass fraction can be established: if the nutrient uptake enzyme has an exceptionally high catalytic rate, its proteome fraction nearly vanishes. Then, metabolite cost 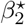 nearly vanishes, and 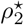 can reach arbitrary values, which on their own may exceed 2*𝓁* drastically.

#### Comparison to experimental data

We compute several observables regarding the whole cell under the assumption of optimal resource allocation in our model. For physiological kinetic parameter distributions, they are in good agreement with those observed empirically (Fig. S2), apart from a slight overestimation of the cellular metabolite mass fraction. The metabolic resource allocation patterns identified here are generic and independent of the mean of *κ*^max^. They are also largely independent of the variance in *κ*^*max*^, as long as individual enzymes do not dominate the proteome. Overall higher efficiencies solely lead to higher growth rates and fluxes. Higher values of *K*, on the other hand, lead to higher dilution relative to growth, creating more significant flux gradients. This would also result in stronger position-dependent gradients in optimal metabolite saturation.

### Optimal balanced growth states under nutrient limitation

We consider an additional constraint on the nutrient uptake flux from the environment (which was left unconstrained in the nutrient-rich regime, and jointly optimized with the rest of the metabolic state of the cell), next to the constraint on the cellular protein concentration. We then identify the metabolic state, which maximizes growth rate at the given nutrient uptake flux. In the absence of such a constraint on nutrient uptake flux at low nutrient concentrations, the complete calculus presented above would remain valid at low nutrient concentrations. Metabolite production costs for all intracellular metabolites would remain on the order of 1, and all *ϕ*_*i>2*_ would converge to zero with decreasing flux *j*_1_ according to Eqs. [**12**], [**11**] and *j*_*i*_ *< j*_1_. This implies that the uptake protein would become a macroscopic fraction of the total proteome, and eventually the whole proteome. Instead, for the reasons outlined in the main text, we specifically assume that uptake protein expression is constrained, and thereby limits the nutrient uptake flux. Consider any constraint on *ϕ*_1_(*ρ*_1_). To find optimal resource allocation under the constraint, we use the method of Lagrange multipliers,

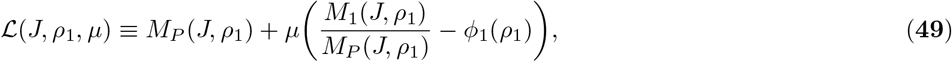

with a Lagrange multiplier *µ. The op*timal resource allocation must now fulfill 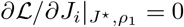 for *i* = 1, …, *𝓁* − *1, in* analogy to Eq. [**34**], and additionally *@* /*@µ* = *0. For i 2 the up*take mass *M*_1_ *does n*ot depend on *J*_*i*_, and the partial derivatives are

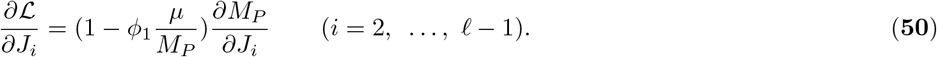

They are zero in the optimal state if Eq. [**5**] is fulfilled. Thus the iterative computation of optimal fluxes [**37**] is still valid. *J*^⋆^ and all 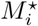 for *i in* (2, …, *𝓁*) follow from the iterative computation of optimal fluxes. A different boundary condition *ρ*_1_ now matches a certain starting condition of the iteration 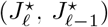 in comparison to unconstrained optimality. For a given constraint *ϕ*_1_(*ρ*_1_), this boundary condition can be found by solving

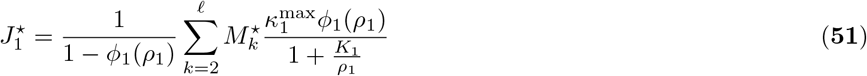

for *ρ*_1_, which corresponds to solving *∂ℒ*/ *∂µ* = *0. The* last remaining condition on optimality is 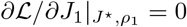, which could be solved for the remaining undetermined variable *µ. Howeve*r, *µ is not r*equired for describing the optimal state, and equations given above [**37, 51**] fully suffice to compute optimal balanced growth states under nutrient limitation. The iterative construction of optimal balanced growth states is applicable under nutrient uptake constraints.

#### Metabolite costs under nutrient limitation

To put this result into the context of metabolite costs, we again consider the variation of a single metabolite concentration (and drop the superscripts ·^⋆^ below). Based on Eq. [**4**], we derive

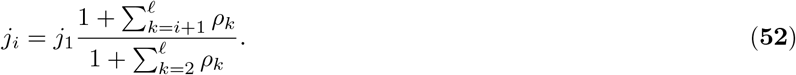

This means that under a constraint on nutrient uptake, which defines a fixed flux *j*_1_(*ρ*_1_) in a given nutrient environment, variation of a single metabolite concentration violates growth optimality, *∂j*_*𝓁*_/*∂ρ*_*k*_ ≠ 0. A variation of a single metabolite concentration that maintains all other metabolite concentrations is ill-defined. Instead, we reformulate the optimality condition. We observe that in an optimal balanced growth state under a nutrient uptake constraint, the sum of metabolite concentrations must be minimal to maximize growth [**52**]. This criterion can be reformulated further: all variations of any single metabolite concentration must have the same effect on the sum of proteome mass fractions 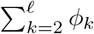

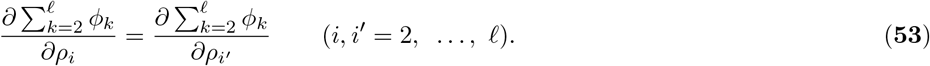

To verify, suppose that this statement was incorrect in a growth-optimal state. A joint variation of *ρ*_*i*_ and *ρ*_*i*_*’* by the same *δ*ρ* in oppos*ite directions could reduce the required sum of proteome fractions. Adding the freed proteome fraction to any reaction would reduce dilution fluxes and thereby improve growth rate.

We insert the kinetic growth law 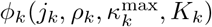, and find

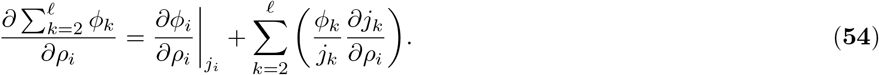

We insert [**52**] and subtract Eq. [**53**] for *I*^*’*^ = 2,

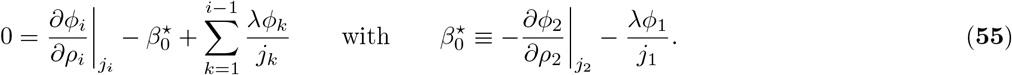

A single summand is added in comparison to the unconstrained optimality condition [**40**]. This additional cost associated with nutrient uptake affects all metabolite production costs equally as an additive constant. We obtain the functional form 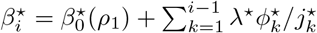, which is Eq. [**10**].

Again, local metabolite production costs are not suitable for determining optimal resource allocation, as they depend on the optimal resource allocation towards other reactions. Instead, we use them to interpret patterns in optimal resource allocation. We therefore do not solve for 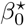 analytically, but simply identify the additional summand as an arbitrary offset, which depends on the nutrient environment and the nutrient uptake constraint. Under variation of *ρ*_1_ we biologically expect to find a one-parametric family of optimal states that fulfill the constraint.

Instead of characterizing optimal states by their nutrient concentration *ρ*_1_, we choose to characterize them by their growth rate 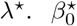 is an arbitrary function of *λ*^⋆^ in this scenario. We can analogously to the case of unconstrained optimality derive 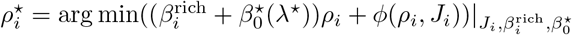. Here, 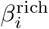 denotes the expression found for 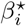 in the nutrientrich regime 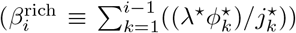. The shape of the equation, including an additive 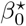 is independent of the exact nature of the constraint limiting nutrient uptake, as long as it only affects the flux *j*_1_. *Notab*ly, a diffusive constraint on nutrient uptake leads to the same result: in this case, the mass required for increasing the nutrient uptake flux *j*_1_ *would* be larger than *δj*_1_/*κ*_1_, because the local nutrient concentration at the cell surface would decrease. This also yields an additional cost 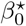, which affects all reactions equally. Based on this argument, we here assume that the additional cost 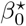 vanishes in a nutrient-rich environment. In a nutrient-rich environment, the balance between diffusion and uptake has a vanishingly small effect on the metabolite concentration at the cell surface. Additionally, uptake protein expression was observed to increase under nutrient limitation, making it unlikely that the uptake protein expression is constrained below its unconstrained-optimal value in nutrient-rich environments. The effect of a regulatory nutrient uptake constraint on uptake expression, growth and metabolite cost is illustrated in Fig. S3.

#### The limit of vanishing nutrient uptake

Above, we derived that a nutrient uptake limitation affects all reactions equally by an additive cost component 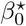. Solving for 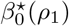 for a given uptake constraint *ϕ*_1_(*ρ*_1_) is challenging due to the same feedback loops concerning optimal balanced growth states discussed earlier. However its form as an additive constant allows for a simplified computation based on the summation constraint on proteome fractions,

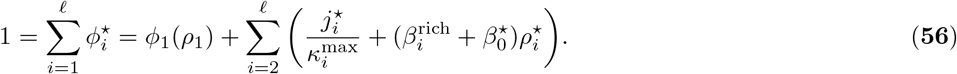

In the limit of strong nutrient limitation *j*_1_ *→*0, this constraint can be solved directly for 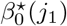: By Eq. [**4**], all fluxes fulfill *j*_*i*_ *< j*_1_ and vanish in the limit. Thus, each *ϕ*_*i*_ *is wel*l approximated by the linear term 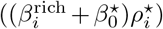 of Eq. [**12**]. We denote 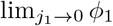 with 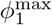. By inserting Eqs. [**11**] and [**10**], we find

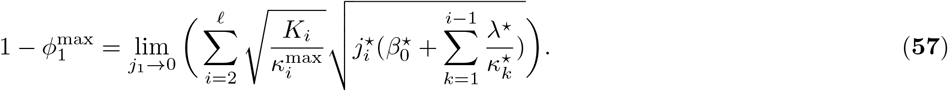

For the reasons outlined in the main text, we assume that the uptake enzyme never constitutes the whole biosynthetic proteome, even in the limit of vanishing *j*_1_, *and* 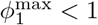. Thus, the right-hand-side must also remain finite in the limit. For a given *j*_1_, growth is maximized if the metabolite mass fraction *M*_*R*_/*M*_*P*_ *is* minimal [**52**]. We show below by construction, that *M*_*R*_/*M*_*P*_ *→* 0 in the limit of *j*_1_ *→* 0. We expand in small *j*_1_ *and dr*op all terms of order *j*_1_. *We sp*ecifically use 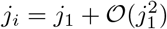, and that all terms 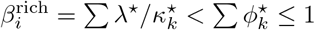 remain finite in the limit,

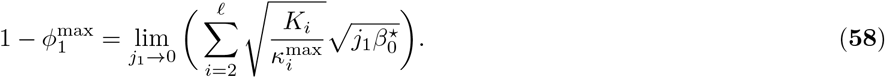

Under a strong constraint on nutrient uptake, 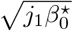 must therefore converge to a suitable normalization constant *Z, and* 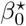 must diverge as 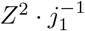. This means that resource allocation in the limit of vanishing nutrient uptake is given by

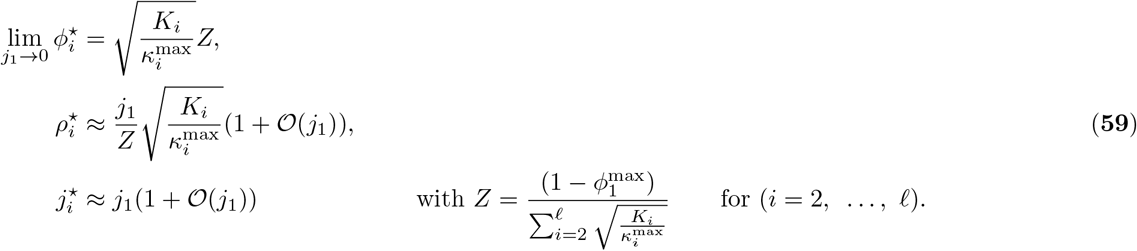

As assumed above, metabolite concentrations vanish in the limit of vanishing nutrient uptake. In this limit, optimal proteome fractions are not anymore set by enzyme efficiencies, but by their specificities.

#### Expansion to small nutrient uptake flux

An optimal balanced growth state in the limit of vanishing nutrient uptake is experimentally unattainable. For this reason, we compute optimal balanced growth states at small but finite growth rates by a first-order expansion of Eq. [**56**] in small *j*_1_ = *λ*^⋆^(1 + 𝒪 (*𝓁*)),

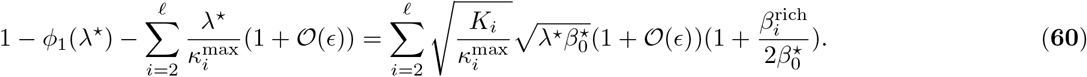

As long as 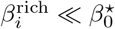, we obtain

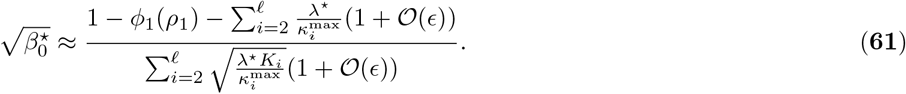

We specifically find for *ρ*_*i*_ by Eq. [**11**], and for *ϕ*_*i*_ *by Eq*. [**56**],

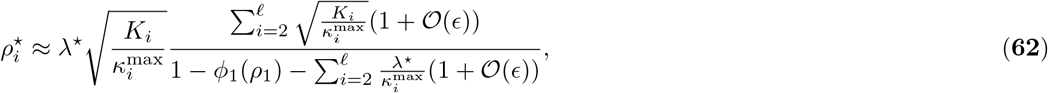

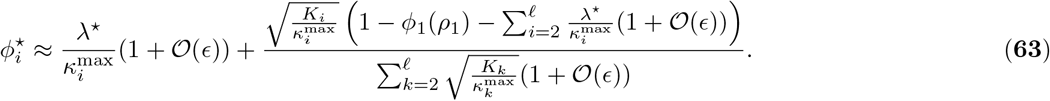

Proteome mass fractions are approximately linear in growth rate in the limit of strong nutrient limitation (apart from nonlinearities introduced by *ϕ*_1_(*ρ*_1_), and metabolite dilution terms of order *ϵ*) and converge to well-defined values as derived above. Correction terms are of order 𝒪 (*ϵ*) in Eqs. [**62**] and [**63**]. For physiological parameter values, they can thus be neglected even at higher growth rates, and the approximation above remains valid as long as 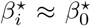. For this, 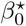 must be larger than all 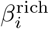. We estimate based on Eq. [**61**], and the definition of 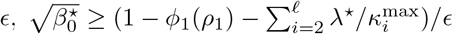 Position-dependent metabolite production costs 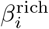 do not exceed 1. If the constraint on *ϕ*_1_ is stringent enough to. ensure 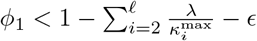, the added cost exceeds 1. Thus, proteome fractions are approximately linear in growth rate at growth rates

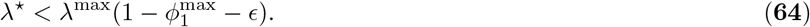

This constitutes the boundary between the nutrient-rich and the nutrient-poor regime. The boundary translates approximately into *ρ*_1_/*K*_1_ *≃* 1: Above the boundary, the catalyzed flux through the first reaction is largely unaffected by reductions of *ρ*_1_, and growth rate is approximately *λ*^*max*^. Below 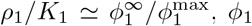 is not up-regulated meaningfully further, a property discussed in more detail in the SI section on regulatory constraints. Then, any further reduction in *ρ*_1_ will lead to a near-proportional reduction in *λ*, and *β*_0_ quickly exceeds one. The approximate equivalence of 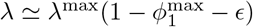 with *ρ*_1_/*K*_1_ *≃* 1 was further verified numerically (Fig. S2F).

#### Linearity of expression level changes

Under physiological conditions, both 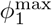 and *ϵ* are typically small. Neglecting both, an even stronger approximation can be made to simplify proteome fractions as a function of growth rate [**63**],

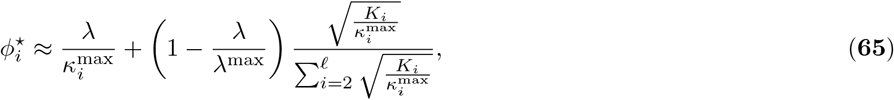

which is a linear function in growth rate for each enzyme. We normalize by the expression level in the nutrient-rich regime 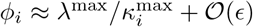. Inserting the definition of *q* and the approximate maximal growth rate for negligible *ϕ*_1_ and 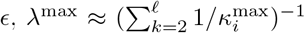, we find that an enzyme’s biochemical quality directly determines the optimal expression fold-change between a nutrient-rich environment and the limit of vanishing nutrient availability,

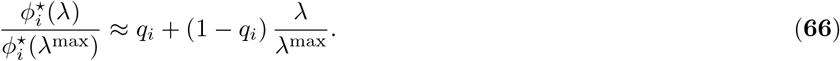

In a chain of irreversible Michaelis-Menten reactions with physiological parameters, any constraint that restricts the uptake proteome fraction from becoming a macroscopic fraction of the proteome under nutrient limitation leads to an optimal resource allocation pattern approximately linear in growth rate for all enzymes except the uptake enzyme. Enzymes with *q* < 1 are down-regulated upon nutrient limitation, enzymes with *q* > 1 are up-regulated. If a finite *ϕ*_1_ is to be taken into account, the critical value of *q that dif*ferentiates between up- and down-regulation shifts to 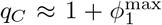. Further non-linear corrections arise at high growth rates due to finite *ϵ* and, in particular at very high *q*, due to the lowered substrate saturation in the nutrient-rich regime (Fig. 2). However, these corrections are typically small in the physiological parameter regime (Fig. 4).

### Model evaluation against physiological cell characteristics

#### Applicability to individual cascades

Our calculations indicate that the discovered two-dimensional maps, flux- and saturation gradients as well as linear expression level changes upon nutrient limitation are not just properties of averages, but can be recovered for individual cascades. We illustrate this result in Fig. S4. Quantitatively, position and biochemical quality account for about 98 % of the variances in scaled metabolite concentrations and expression levels respectively in the nutrient-rich regime, as well as about 80 % of the corresponding variances under nutrient limitation in our ensemble of optimal resource allocation in physiologically parameterized cascades. These fractions denotes percent reductions in squared residuals before and after applying the re-scaling derived through our analytical calculations (see Fig. S4). We thus expect our results to be applicable to a single physiological parameterization, as would be encountered in a given experiment.

#### Predicted correlations in proteomics and metabolomics datasets

Our model predicts a correlation of protein expression fold-changes and protein expression levels, as both depend on biochemistry. For physiologically small dilution strength, most enzymes are well-saturated, and the quadratic term dominates Eq. [**12**],

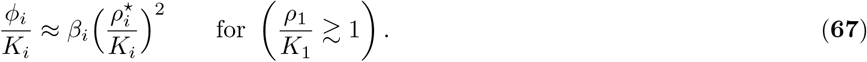

We insert Eq. [**11**] and *j*_*i*_ *≈ λ* + *𝒪* (*ϵ*), to find 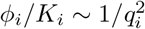. Together with Eq. [**66**], this defines the correlation

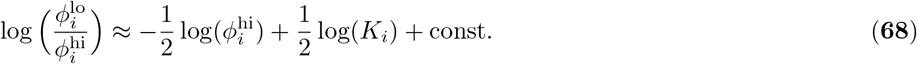

We predict an anti-correlation with log-log-slope of -1/2 between expression levels in the nutrient-rich regime and expression fold-changes in the limit of vanishing growth. We note that the affinity-dependent term log(*K*_*i*_)/2 may introduce considerable noise. If no affinities are known, averaging over a large number of enzymes may be required to identify the anti-correlation. We here test whether an anti-correlation is present when taking the entire proteome into account (Fig. 5). We note that this approach is biased for several reasons: the proteome contains not only enzymes involved in the conversion of environmental nutrients into new cell biomass, but for example also structural proteins and stress-response proteins. Our model does not account for such other proteome constituents, and does not predict their growth responses. At the same time, our model suggests that protein complexes should be treated as a joint proteome fraction. Still, we observe an anti-correlation stronger than expected from regression to the mean, both in the entire proteome, and when restricting our analysis to individual metabolic pathways, which consist solely of proteins involved in biomass synthesis.

Our analysis of the proteome could be improved by including additional kinetic or metabolomic data. Unfortunately, accurate metabolomic and kinetic data is not available for many reactions. A small number of reactions was recently analyzed by Dourado et al. (16). In this dataset, quantitative deviations from the original enzyme-substrate relationship appear too small for the considered enzymes for meaningful analysis. We expect that ongoing efforts to expand the availability and accuracy of kinetic data (11) will allow for a direct test of our predictions. Under nutrient limitation, we predict that the added cost of metabolite production should shift the optimal enzyme-substrate relationship to lowered metabolite concentrations. Interestingly, a qualitatively similar deviation from the un-shifted enzyme-substrate relation was observed experimentally when reducing growth rate by switching the carbon source of a growth medium (16).

### Extension to more complex network topologies

Metabolic networks typically exhibit a complex topology. Capturing the full complexity of a real cell lies beyond the scope of this paper, as it would impede analytical solvability, vastly increase the degrees of freedom in the parameterization, and impede the identification of simple patterns. However, we use an exemplary network with more complex topology to illustrate that our framework applies to more complex topologies and that its key predictions are still valid. We focus on one particularly interesting use case that can be directly linked to existing proteomics studies: the consumption of two different, non-interchangeable environmental nutrients by the cell, such as a carbon and a nitrogen source. We specifically model this system as two convergent pathways with two different environmental nutrients (Fig. 6E). To ease the notation, we call one pathway the N-branch and the other the C-branch. At the convergence point, both metabolite types (C-type and N-type derived from the two nutrients) are required for a single reaction. The product of this reaction is further converted by a third pathway, the stem of the metabolic network.

We assume that the enzyme at the convergence point exhibits Michaelis-Menten-kinetics in both of its substrates - i.e. both substrates are needed for the reaction to proceed,

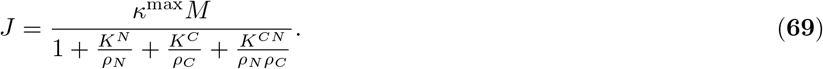

This rate law features a kinetic constant *K*^*CN*^ (with units of concentration squared), for which no physiological parameter regime is known. We here assume that *K*^*CN*^ = *K*^*N*^ *K*^*C*^. We further extend the framework of mass conservation to the convergence point: we assume that the two metabolite types are used with a fixed mass stoichiometry. The C-branch contributes a mass flux fraction *γ*_*C*_ *to the* downstream mass flux, the N-branch *γ*_*N*_ =*1 γ*_*C*_, with 0 *< γ*_*C*_ *< 1*.

Enzymes along the C-branch and the stem form a chain of enzymes. Its enzymes are numbered as before. On the other hand, enzymes in the N-branch are numbered by their position in the branch. To differentiate, we denote indices in the N-branch by a tilde 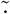. The N-branch has length 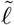. Lety be the index of the enzyme with two substrates, so that *ρ*_*C*_ *ρ*_*y*_. Mass conservation demands

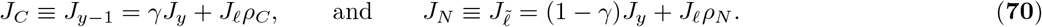

We can now express *M*_*y*_ as a function of only fluxes and kinetic constants,

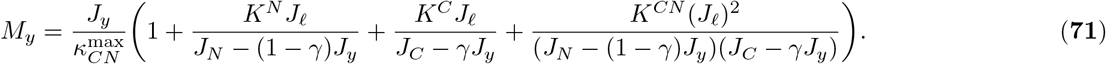

All other enzyme masses are still given by Eq. [**30**]. Thus, all fluxes from 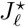 to 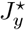 can still be constructed iteratively according to Eq. [**37**] from a starting condition of the iteration 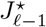 (and 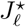, which sets system size). Analogously, we can pick another starting condition 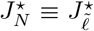 for the N-branch of the cell. This starting condition reflects an additional degree of freedom in space of growth-maximizing metabolic states solution: next to system size (associated with *J*_*𝓁*_) and nutrient environment (associated with *J*_*𝓁*−1_), which were both also present in the linear chain, an optimal state is now also dependent on a second nutrient concentration in the environment (associated with *J*_*N*_). For any *J*_*N*_, one can solve *∂M*_*P*_/*∂J*_*y*_ = 0 for *J*_*C*_ analytically by a suitable computer algebra system.

For all other criteria on optimality, Eq. [**71**] behaves like a regular Michaelis-Menten kinetic rate law with

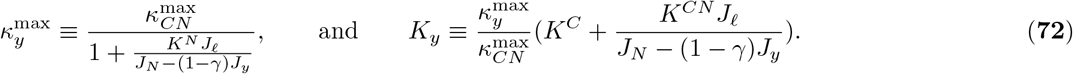

Thus, the iterative construction of the optimal metabolic state can proceed until 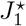, where a matching environmental nutrient concentration must be determined. In comparison to the solution of a linear chain with *𝓁* enzymes, all fluxes *J*_*i<y*_ are scaled by *γ*. Analogously, all fluxes in the N-branch can be constructed, once 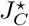 is known by viewing the N-branch and the stem together as a linear chain. In this manner, a consistent solution is always found, because the same equation *∂M*_*P*_/*∂J*_*y*_ = 0 is used to determine first *J*_*C*_ from a given *J*_*N*_ and then *J*_*N*_ again from the determined *J*_*C*_. Due to the irreversible nature of reactions considered, matching environmental nutrient concentrations *ρ*_1_ and 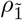 can be determined (even for constrained expression levels *ϕ*_1_ and/or 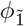), analogously to the linear chain. To parameterize the branched network, we assume that the substrate mass *M*_*S*_ on each branch is given by *γ* times the substrate mass present in the stem of the network. While *M*_*E*_, *κ*, and the distributions of *k*_*cat*_ *and* 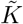 are the same as in the stem, thus both *K* and κ^*max*^ are on average reduced by a factor of *γ* in comparison to their values in the stem.

In the nutrient-rich regime, optimal resource allocation in two convergent pathways qualitatively resembles optimal resource allocation in a single chain. Scaled metabolite concentration gradients in each separate pathway are qualitatively unaffected by the altered topology. They exhibit a strong decrease over the first few reactions after nutrient uptake and a weaker decline after. At the convergence enzyme itself, the cost of metabolite production suddenly increases - requiring only one production pathway beforehand, but both in the shared stem. Thus, saturation drops at the convergence.

If one of the two nutrients is available at a growth-limiting concentration, the proteome reallocation within the branch with limited nutrient influx and in the stem resembles the proteome reallocation in a single chain: high-*q* enzymes are up-regulated while low-*q* enzymes are down-regulated (Fig. 4D, F). On the other hand, proteome fractions in the unaffected branch are reduced with decreasing growth rate, leading to an overall resource reallocation from the unaffected branch and low-*q* enzymes in the affected branch and stem to high-*q* enzymes in the affected branch and stem.

### Regulatory constraints

#### Regulation of the uptake enzyme

In a simple regulatory model, a metabolite can directly act as a transcription factor to influence protein expression. Here we assume that such a regulatory framework leads to a Hill-function governing an enzyme’s expression level as a function of its substrate’s concentration, with a basal expression level *ϕ*^0^ and an expression level *ϕ*^*∞*^ at full promoter saturation with the metabolite. At a concentration *K*^*reg*^, the promoter is half-saturated,

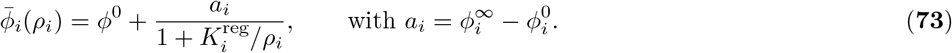

In this model, the metabolite can either induce protein production (*ϕ*^0^ < *ϕ*^*∞*^), or inhibit protein production (*ϕ*^0^ *> ϕ*^*∞*^). We focus first on the nutrient uptake reaction (*i* = 1). *Co*nsider a cell that efficiently converts environmental nutrients into new protein biomass (*J*_1_ *≈ J*_*𝓁*_ (1 + 𝒪 (*ϵ*))) and that exhibits a small uptake protein mass fraction *ϵ*_1_ *≪ 1 in a* nutrient-rich environment. The flux that can be efficiently handled by downstream metabolism (*i* = 2, …, *𝒪*) is proportional to 1 _−_ *ϵ*_1_. For sufficiently small *ϕ*_1_ and only moderate reductions in nutrient availability, keeping the nutrient uptake rate approximately constant therefore recapitulates the unconstrained optimal resource allocation strategy,

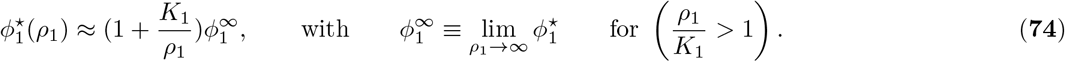

In the case of the nutrient uptake reaction, we thus expect that the proteome fraction of the uptake enzyme should increase with decreasing nutrient availability (Fig. S3A). For the reasons outlined in the main text, we assume that the approximate form [**74**] is only valid at high growth rates. Instead, expression levels off to 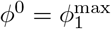 well before the uptake protein becomes the dominant proteome mass fraction in the cell. To achieve qualitatively comparable up-regulation of all parameterizations in the ensemble despite different optimal expression levels in a nutrient-rich environment, we use 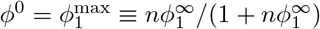, where n determines the fold-change of *ϕ*_1_ under nutrient limitation. In this manner, *ϕ*_1_ can be up-regulated n-fold for parameterizations with small *ϕ*^*∞*^ but does not exceed 1 for larger *ϕ*^*∞*^. In all simulations, we use *n* = 5. To mimic optimal resource allocation [**74**], *K*^*reg*^ *must* then be 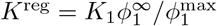. This regulatory scheme approximates unconstrained optimal resource allocation to the uptake protein at high nutrient concentrations *ρ*_1_/*K*_1_ *»* 1, and smoothly interpolates to a cellular state with a constrained nutrient uptake flux at lower nutrient concentrations (Fig. S3).

#### Approximating optimal resource allocation by regulation

We define an extended regulatory scheme for system-wide metabolism. Transcription takes place with enzyme-specific rates

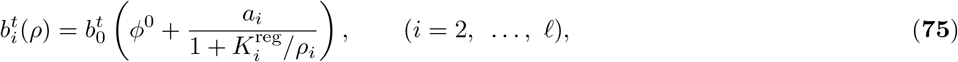

where 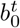 is a basal transcription rate. We use metabolite-mediated regulation as introduced in the main text, which is specified by the set of parameters ℛ_*m*_(*κ*^*max*^,*K*) = (*ϕ*_0_, *a, K*^reg^)( *κ*^*max*^,*K*) with

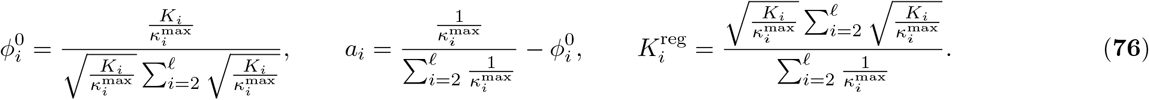

As a second step, we use a minimal model of translation that specifies protein synthesis rates proportional to the corresponding transcription rates,

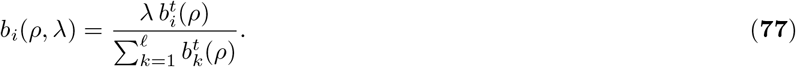

This form sets the overall normalization of protein synthesis, Σ_*k*_ *b*_*i*_(*ρ, λ*) = *λ*, as required in a steady state with dilution at rate *λ*, and leads to stationary expression levels 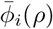 given by equation [**24**]. The relation between transcription and translation rates reflects competition for a growth-dependent global resource, e.g., RNA polymerases, translating ribosomes, or energy.

The steady-state protein expression levels given by the metabolite-mediated regulatory scheme [**75** – **77**] reproduce growth-optimal metabolic states in very good approximation. First, we obtain the consistency relation

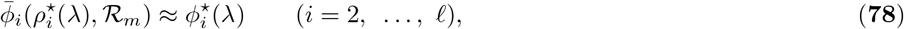

where 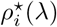 are 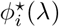 given by equations [**19** – **21**]. Second, we numerically study time-dependent metabolic states 𝒮 (*t*) = (*ρ*(*t*), *ϕ* (*t*)) given by the dynamical equations

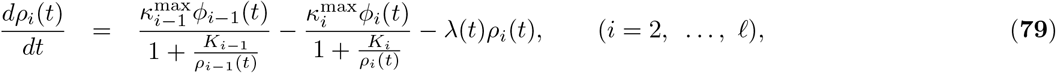

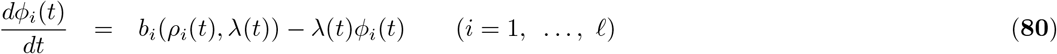

with 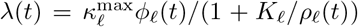 and a time-dependent input signal *ρ*_1_(*t*) (Fig. 6). We find stable stationary states 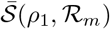 with

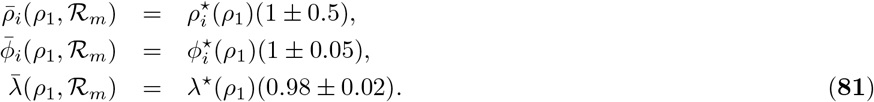

We note that no flux-sensing or growth-sensing mechanism is required to robustly implement growth-dependent gene expression.

### Network evolution

In this subsection^*^, we estimate selection on enzyme-metabolite binding in networks with a regulated physiological response specified by a regulation function depending on the reaction parameters, ℛ (*κ*^*max*^,*K*). We evaluate the selection coefficients, or relative growth effects in the stationary state, of amino acid changes that affect the metabolite binding affinity, *K*_*i*_ *→ K*_*i*_+,*Δ K*_*i*_ (*i* 6 = 1),

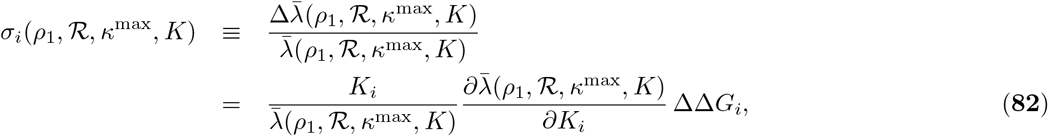

where, Δ., Δ.*G*_*i*_ =, Δ.*K*_*i*_/*K*_*i*_ is the corresponding change in reduced free energy of binding. We assume uniform effects, |Δ., Δ.*G*_*i*_| = 1, in line with typical amplitudes of mutations affecting protein evolution. In equation [**82**], we compute only the direct effect of the mutation on binding, while keeping the regulation function 𝒮 (*κ*^*max*^,*K*) at the wild-type parameters. We assume that regulation tunes the wild-type metabolic state 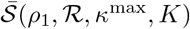 to near-optimal growth,

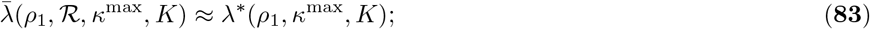

an example of such regulation functions is the metabolite-mediated scheme 𝒮_*m*_ discussed above. This assumption is self-consistent if the evolution of protein binding affinities is followed by compensatory evolution of regulation, 𝒮 (*κ*^*max*^,*K*) 𝒮 (*κ, K* +, Δ.*K*). However, details of the regulation function and its evolution are not relevant for our current purpose, to obtain an order-of-magnitude estimate of selection on protein binding affinities.

First, we compute an upper bound for the selection coefficients *σ* _*i*_(*ρ*_1_,, *𝒮 κ*^*max*^,*K*) from a hypothetical system where the mutant is constrained to the same protein expression levels as the wild-type. For this system, we obtain

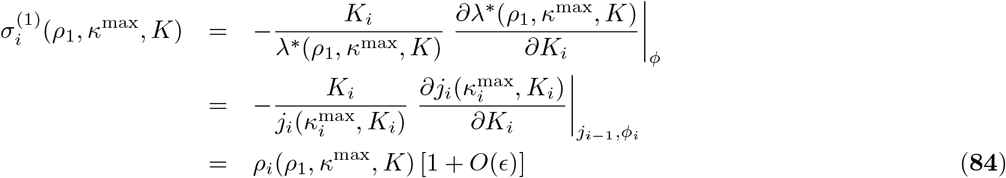

which follows from the Michaelis-Menten relation [**1**], the flux continuity relation [**3**] and 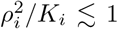 [**11**]. We have used, Δ., Δ.*G*_*i*_ = −1 for an affinity-increasing mutation and the relation [**83**] for the wild-type. We can compare this strength of selection with the case of a single enzyme at constant metabolite density,

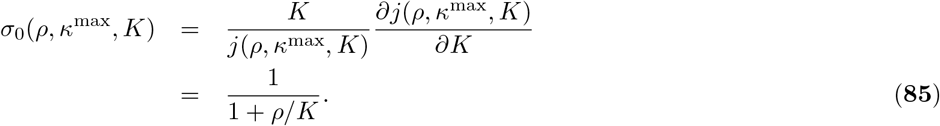

This shows that in a network, moderate affinity changes of bulk enzymes are largely buffered by compensatory changes of the cognate metabolite,, Δ.*ρ*_*i*_/*ρ*_*i*_ −*δK*_*i*_/*K*_*i*_, leaving a residual effect on growth by metabolite dilution. Furthermore, the effects on downstream metabolites, which are omitted in [**84**], are of order *ϵ*^2^.

*Secon*d, we compute a lower bound on *σ*_*i*_(*ρ*_1_,, *κ*^*max*^,*K*) from the variation of optimal growth rates. This approximation couples local enzyme changes *K*_*i*_ *→K*_*i*_ +, 6.*K*_*i*_ and compensatory changes of regulation, 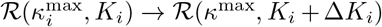, assuming a hypothetical simultaneous evolution of an enzyme’s protein and cis-regulatory sequence, which overestimates the mutant fitness. We obtain selection coefficients

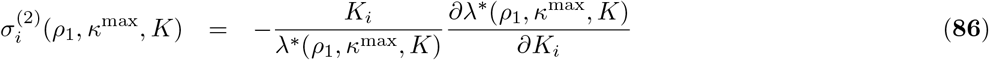

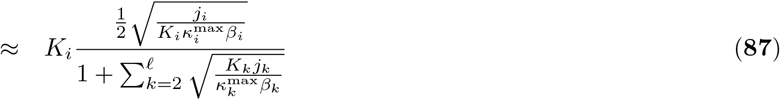

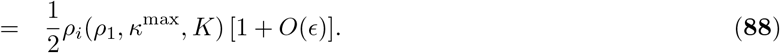

Here we have used that *j*_1_ is independent of *K*_*i*_ (which is justified under nutrient limitation, see equation [**73**], and correct to order *ϵ* otherwise), and neglected position-dependent inhomogeneities and changes of the cost factors (i.e., *β*_*i≈*_ *β*_0_ *≈* const., equation [**21**]), as appropriate under nutrient limitation. In the same rationale, variation of the growth-optimal metabolic state S^*\**^(*ρ*_1_, *κ*^*max*^,*K*) provides estimates of the buffering in regulated networks,

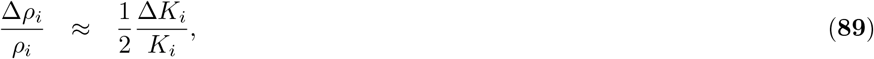

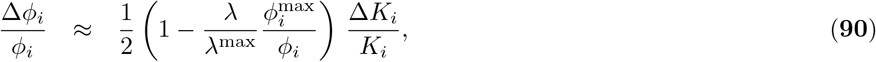

with 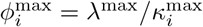, which are obtained from equations [**19**] and [**20**].

The approximations [**84**] and [**88**] of the selection coefficients *σ*_*i*_(*ρ*_1_, *ℛ, κ*^*max*^,*K*) are of the same form and magnitude, which establishes equation [**25**] and the resulting mean field-estimates of *σ*_*i*_ derived in the main text. Furthermore, inserting the metabolite densities given by equation [**13**] or [**19**] shows that individual enzymes evolve target binding on exponential fitness landscapes,

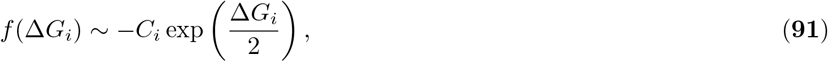

where, Δ.*G*_*i*_ = *log*(*K*_*i*_/*K*_0_) is the reduced free energy of binding, *f*(,*Δ*.*G*_*i*_) = *λ* (, *Δ*.*G*_)_ - λ (,*ℛ*.*G*_*i,min*_) is the growth difference to the protein state of strongest binding, and *C*_*i*_ is a constant. This form is similar to the biophysical fitness landscapes of viable proteins discussed in ref. (58).

### Numerical simulations

Numerical computations were performed using Python. Equations [**6**], [**3**] and [**1**] were used to construct optimal balanced growth states of stochastically parameterized metabolic models from iteration starting conditions (*J*_*𝓁-*1_, *J*_*𝓁*_). Iteration starting conditions were varied numerically using the Broyden–Fletcher–Goldfarb–Shanno algorithm to match a nutrient-rich environment (Fig. 1 – 4, S2 – S4), or match an otherwise defined environmental nutrient concentration (Fig. 3, orange points). Subsequently, the ratio *J*_*𝓁-*1_/*J*_*𝓁*_ was reduced to construct optimal balanced growth states under nutrient limitation. In case of the branched metabolic network, also the metabolite concentration *ρ*_*N*_ was varied numerically for each ratio *J*_*𝓁-*1_/*J*_*𝓁*_ using the same algorithm to ensure a constant environmental nutrient concentration 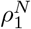 under nutrient limitation of the *C* branch. Time evolutions of the feed-forward regulated metabolic cascades (Fig. 6, Eqs. [**79**] and [**80**]) were computed by means of Euler’s method. The code required to construct optimal balanced growth states and all figures displayed in the manuscript is available online in the following repository: https://github.com/laseeg/ResourceAllocation.

### Cytoplasmic density affects optimal resource allocation

Throughout the whole paper, we assume that the protein mass density of the cell is approximately 400 g L^-1^, independent of environmental conditions. This assumption is in line with experimental observations (1, 26–28), which indicate an approximately constant cellular dry mass density under nutrient limitation. However, literature estimates of the cellular protein mass density vary, and we can not exclude that the composition of the cell varies under nutrient limitation which may lead to a modified biosynthetic protein density at constant dry mass density, or exclude that cellular dry mass density changes within the margins of experimental accuracy. A recent study has proposed potential underlying mechanisms to a constant intracellular protein density based on a model of cellular self-replication similar to the one we present here (67). Simultaneous optimization of protein density and proteome allocation under a nutrient limitation may be feasible but lies beyond the scope of this paper.

Generally, we predict that cells exhibit higher optimal metabolite mass fractions and lowered mean enzyme saturation at lowered protein mass density in the nutrient-rich regime. The metabolite mass fraction changes in proportion to the squareroot fold change in protein mass density. The same behavior is expected under a nutrient uptake constraint, which concerns *ϕ*_1_. Thus, changes in resource allocation induced by changes in protein mass density are likely negligible in comparison to the predicted changes due to differences in biochemical quality *q* at constant protein mass density. Only a constraint on *j*_1_ (but unconstrained *ϕ*_1_ relative to all other metabolic proteins in the cell) qualitatively yields a different behavior. In this case, both the cellular metabolite mass fraction and mean enzyme saturation would increase in the nutrient-poor regime with decreasing biosynthetic protein mass density.

---

**Figure S1:**
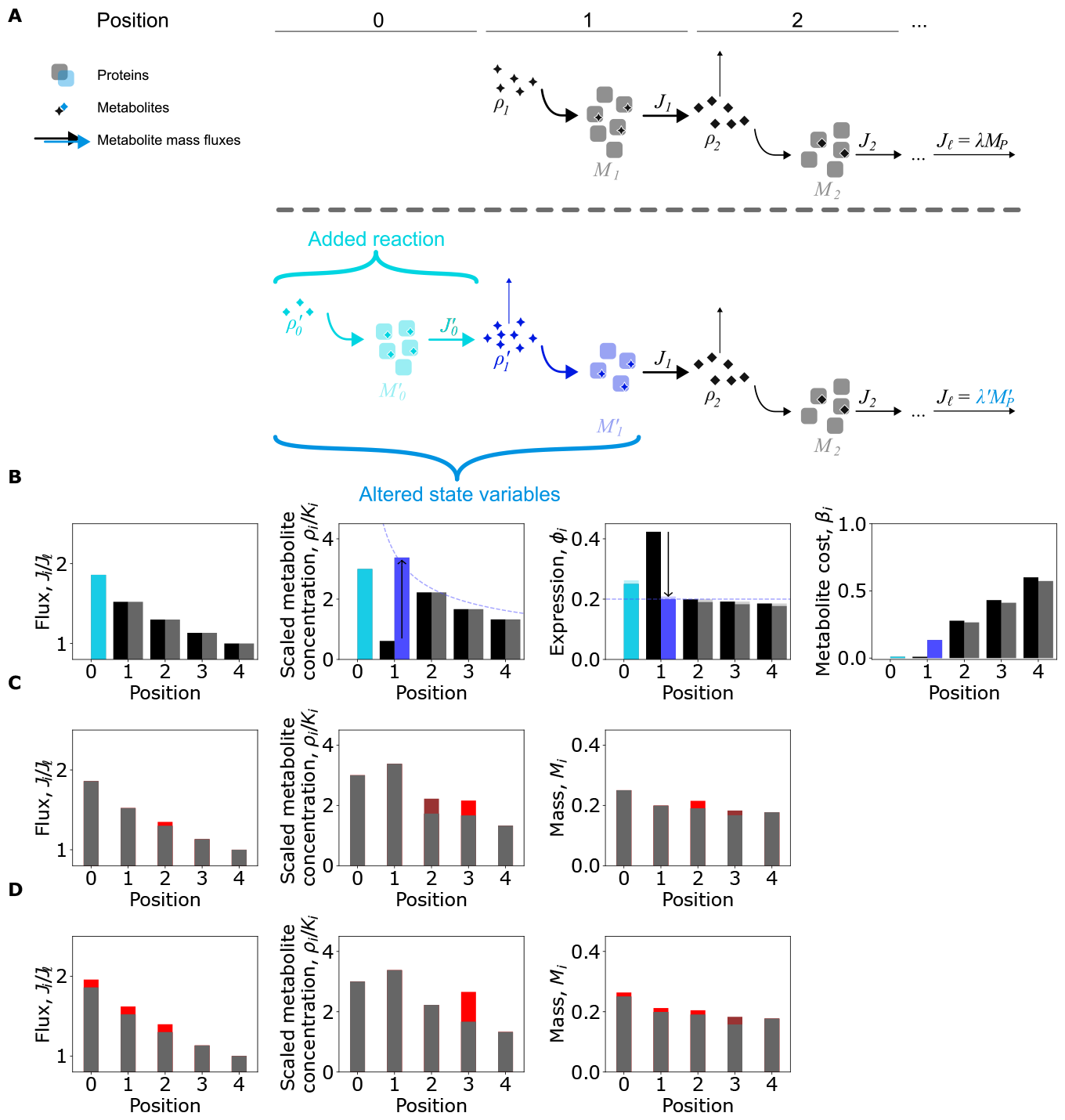
Optimal metabolic states can be constructed iteratively. (*A*) For a two-enzyme chain, the optimal metabolic state in a given nutrient environment *ρ*_1_ can be determined analytically (33). Here we construct optimal metabolic states of longer chains, by iteratively adding an enzyme to the chain that produces the previous environmental nutrient. Suppose all growth-optimal fluxes *J* of a chain of length 𝓁 are known. We show (SI) that a new growth-optimal set of fluxes can be constructed by adding a single new flux 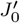 to the chain and finding a suitable concentration of the new environmental nutrient 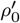. Such an extension of the chain defines a protein mass of the newly added enzyme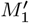, and affects the metabolite mass concentration of the previous environmental nutrient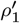, as well as the required mass of the consuming enzyme 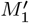, but does not affect further downstream metabolite mass concentrations or enzyme masses, making the problem analytically tractable. (*B*) The optimal metabolic state of an exemplary short chain (left, black) and the optimal state of the corresponding elongated chain (right, shaded) are shown next to each other for comparison. For simplicity, we illustrate a cascade with equal substrate affinities *K*_*i*_ = 0.1 and efficiencies 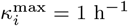 for all enzymes. A system of five equations can be solved for the five modified or new state variables 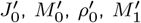 and 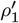. All other fluxes, saturations, and masses are conserved under this elongation. Instead of enzyme masses *M*_*i*_, dimensionless enzyme mass fractions *M*_*i*_/*M*_*P*_ are shown. Light gray bars represent unchanged enzyme masses before rescaling due to the change of 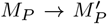, caused by the added mass 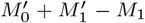. Arrows indicate changes in the metabolite mass concentration 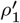 and enzyme mass 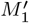 caused by adding a metabolite-producing reaction with index *i* = 0. The resulting patterns in fluxes, saturation, and expression can be understood through the cost of metabolite production *β*_*i*_. The cost reflects how many proteome resources are needed to produce a given metabolite. For physiological chain parameters, the cost of metabolite production is well approximated by the proteome fraction upstream of the reaction of interest. It increases as a function of position. Thus, the optimal metabolite concentration decreases with position. The resulting gradient is well-approximated by (*K*_*i*_)^*-*1/2^ (curved dashed line). At the same time, dilution of these metabolites induces gradients in mass fluxes. Both gradients in flux and enzyme saturation together result in a very weak gradient in enzyme mass fractions along the metabolic chain. Equal allocation to all five enzymes is shown as a horizontal dashed line for comparison. (*C*), (*D*) Changes in metabolic state variables for the perturbation of a growth-optimal state in a single flux or metabolite concentration, as discussed in the main text. (*C*) Upona variation that increases flux *J*_2_ in comparison to the growth-optimal state, the metabolite concentration *ρ*_3_ increases, whereas *ρ*_2_ decreases. At the same time, the enzyme masses required for these fluxes and metabolite concentrations in steady state, change for enzymes 2 and 3. In the optimal state, the total proteome mass must remain unchanged for a small variation in any single flux. (*D*) A concerted variation of all fluxes upstream of metabolite 3 leads to an isolated change in *ρ*_3_. All enzyme masses upstream increase, whereas the mass *M*_3_ is reduced. In the optimal state, the total proteome mass must remain unchanged for a small variation in a single metabolite concentration. In all panels, increases are shown in bright red, whereas decreases are shown in dark red.

**Figure S2:**
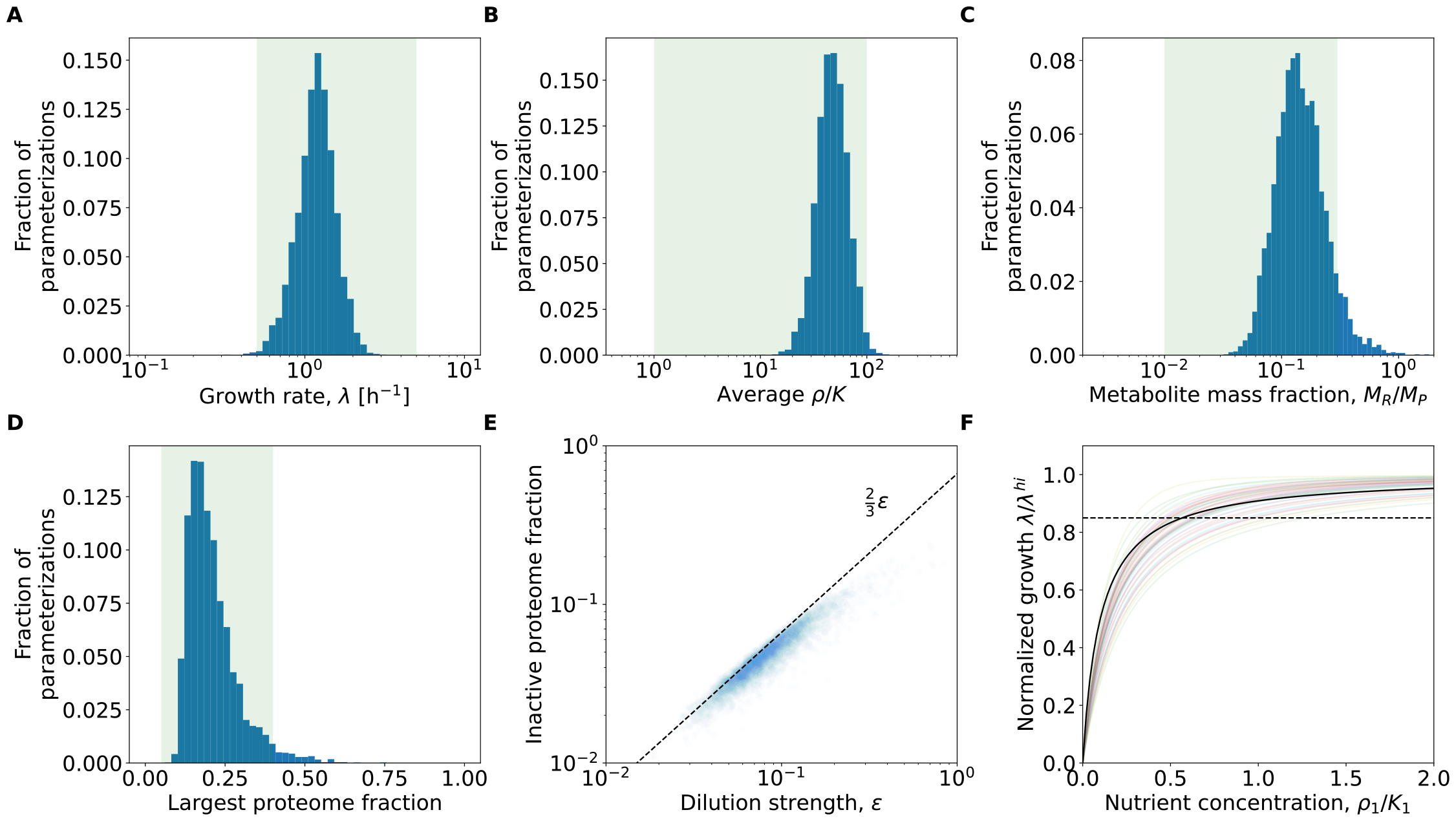
Model cells exhibit physiological properties. An ensemble of 5000 linear chains of enzymes was parameterized by drawing individual enzyme affnities and effciencies from log-normal distributions with physiological mean and variance informed by measurements of Bar-Even et al. (37). The environmental nutrient concentration was set to *ρ*_1_ = 1000*K*_1_ to reflect a nutrient-rich environment. (*A*) Cells parameterized in this manner exhibit physiological growth rates between 0.5 h^*-*1^ and 5 h^*-*1^. (*B*) The geometric mean of scaled intracellular metabolite concentrations (*ρ*_*i*_/*K*_*i*_ for *i* = 2, …, *𝓁*) in physiologically parameterized model cells matches physiological estimates: enzymes are typically operating above their *K*, but not at exceedingly high substrate concentrations. Thus, values between 1 and 100 are deemed physiological. (*C*) Cumulative metabolite mass fraction of all metabolite species in a cell. Typically, metabolites contribute on the order of 10 % of the proteome mass to the total dry mass in a cell (from BNID: 101436). A small fraction of physiologically parameterized cascades exhibit unphysiologically large metabolite mass fractions, exceeding the single percent range. We hypothesize that this result - as well as a relatively high average saturation of enzymes in our model - may stem from neglecting additional growth costs associated with metabolites in a real cell. (*D*) In a typical bacterial cell, the largest single proteome fraction is given by the ribosome and constitutes about 30 % of the cellular proteome. A small fraction of physiologically parameterized cascades exhibit unphysiologically large largest proteome mass fractions. (*E*) Inactive proteome fraction predicted from the simplified resource allocation pattern [**45**] and obtained from optimal balanced growth states in physiologically parameterized cascades in the nutrient-rich regime. (*F*) Predicted, representative nutrient dose-response functions for physiologically parameterized chains with a regulatory constraint on nutrient uptake limiting uptake overexpression to 5-fold (colored thin lines), are well approximated by a Monod-function with *K*_Monod_ = *K*_1_/10 (black solid line). The nutrient-rich regime is separated from the nutrient-poor regime by a growth reduction of 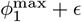 (on average at the black dashed line), which typically corresponds to *ρ*_1_ ≈ *K*_1_.

**Figure S3:**
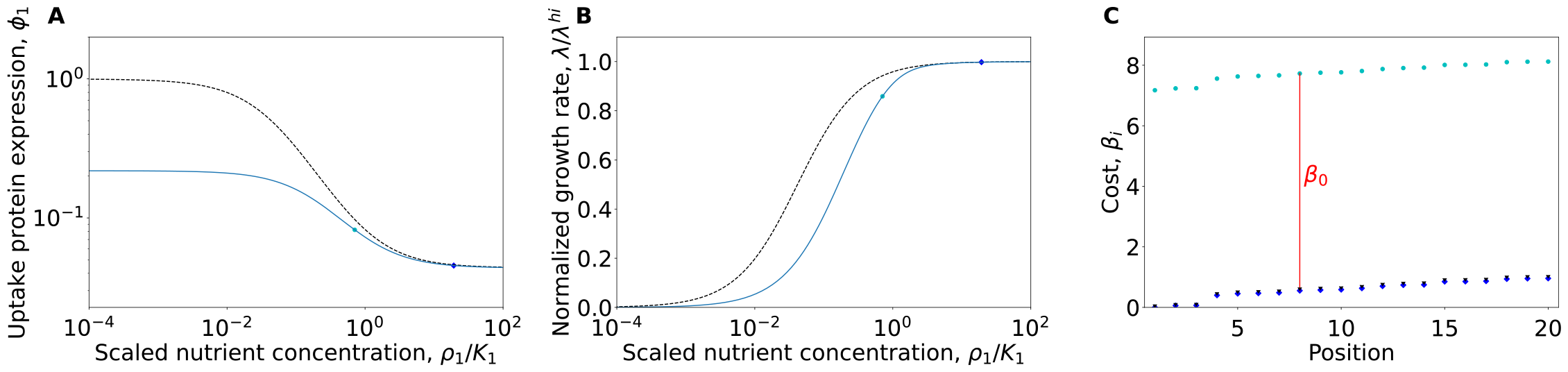
A nutrient uptake constraint increases the costs of metabolite production. (*A*) Unconstrained optimal resource allocation leads to non-physiological overexpression of uptake proteins under nutrient limitation (black, dashed). This ultimately leads to a cell consisting solely of nutrient-uptake enzyme. In contrast, we constrain uptake protein expression (blue) at low nutrient concentrations. The constraint was chosen such that it mimics unconstrained optimal resource allocation at high growth rates and exhibits the same Hill-like shape (SI). (*B*) Growth is reduced in a Monod-like fashion in response to nutrient limitation for both unconstrained (black) and constrained (blue) overexpression of the uptake protein, with a Monod-constant on the order of the nutrient affnity of the uptake protein. The Monod-like growth response results mainly from the Michaelis-Menten kinetics of the nutrient uptake reaction but is modulated by the uptake expression change, as well as the increasing overall effciency of nutrient conversion under nutrient limitation. (*C*) When uptake expression is not constrained, the cost of metabolite production 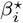 increases several-fold along the metabolic chain (dark blue). It is closely approximated by the cumulative upstream proteome fraction. When the nutrient uptake is reduced below its optimal level, all costs increase (cyan dots). The additional cost is the same for all enzymes 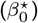. While the cost of metabolite production still increases monotonously, relative changes are reduced. When accounting for differences in *q*, this monotonous increase in cost along the chain leads to a monotonous gradient in metabolite saturations along the chain. As relative gradients in cost flatten upon nutrient limitation, concentration gradients flatten. This leads to a position-dependent fold change in metabolite concentration upon nutrient limitation. All panels illustrate a single, representative, physiological parameterization of a heterogeneous chain of length 𝓁 = 20.

**Figure S4:**
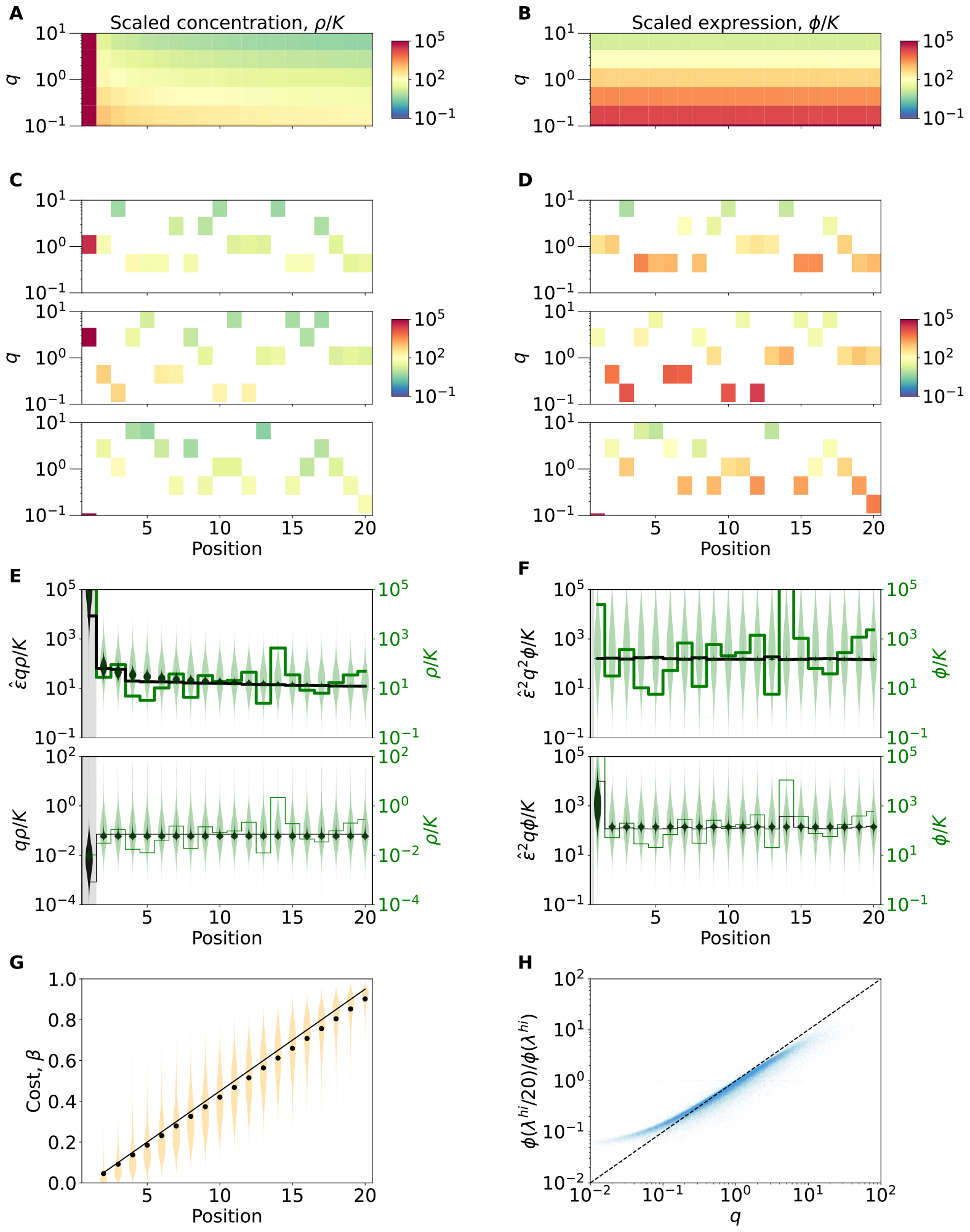
Position and biochemical quality-based predictions are accurate for individual enzymes. (*A*), (*B*) Analytical maps of optimal scaled metabolite concentrations and expression levels. (*C*), (*D*) Corresponding maps of optimal scaled metabolite concentrations and expression levels for three individual model parameterizations. (*E*), (*F*) To quantify the residual variability, unexplained by position and quality, we rescale growth-optimal concentrations and expression levels according to our theory by *q*. The normalization of *q* ensures that the mean value of all *q*_*i*_ is one - this enables a direct comparison of unscaled patterns (green) and scaled patterns (black). Both are shown for a single parameterization of the model cell (shown in the third line of *C, D*, lines). Distributions of scaled metabolite concentrations and expression levels before and after rescaling by *q* and ϵ in nutrient-rich and poor environments from the ensemble of physiologically parameterized chains are shown as violins. To conserve the mean of each distribution and allow for a direct comparison of both scaled and un-scaled variables 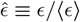 was used for rescaling. Our theory reduces the squared residuals per enzyme of the log concentrations in the nutrient-rich regime by a factor 40, in the nutrient-poor regime by a factor of 6, and the squared residuals of the log expression levels by 50- and 5-fold respectively. (*G*) Distributions of metabolite production costs in an ensemble of physiologically parameterized chains as a function of enzyme position. The solid line corresponds to the estimate of 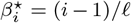. Black dots denote the means of the shown distributions. (*H*) Correlation between enzyme quality and expression fold change upon severe nutrient limitation for all individual enzymes of the ensemble (excluding nutrient uptake enzymes).

## Notes

### Competing Interest Statement

The authors have declared no competing interest.

